# Deep neural network affinity model for BACE inhibitors in D3R Grand Challenge 4

**DOI:** 10.1101/680306

**Authors:** Bo Wang, Ho-Leung Ng

**Affiliations:** Biochemistry & Molecular Biophysics, 141 Chalmers Hall, Kansas State University, 1711 Claflin Rd., Manhattan, 66506, KS, U.S.A.

**Keywords:** D3R, Docking, Binding affinity, Structure-based drug design, Machine learning, Deep neural network, BACE, beta-secretase 1

## Abstract

Drug Design Data Resource (D3R) Grand Challenge 4 (GC4) offered a unique opportunity for designing and testing novel methodology for accurate docking and affinity prediction of ligands in an open and blinded manner. We participated in the beta-secretase 1 (BACE) Subchallenge which is comprised of cross-docking and redocking of 20 macrocyclic ligands to BACE and predicting binding affinity for 154 macrocyclic ligands. For this challenge, we developed machine learning models trained specifically on BACE. We developed a deep neural network (DNN) model that used a combination of both structure and ligand-based features that outperformed simpler machine learning models. According to the results released by D3R, we achieved a Spearman’s rank correlation coefficient of 0.43(7) for predicting the affinity of 154 ligands. We describe the formulation of our machine learning strategy in detail. We compared the performance of DNN with linear regression, random forest, and support vector machines using ligand-based, structure-based, and combining both ligand and structure-based features. We compared different structures for our DNN and found that performance was highly dependent on fine optimization of the L2 regularization hyperparameter, alpha. We also developed a novel metric of ligand three-dimensional similarity inspired by crystallographic difference density maps to match ligands without crystal structures to similar ligands with known crystal structures. This report demonstrates that detailed parameterization, careful data training and implementation, and extensive feature analysis are necessary to obtain strong performance with more complex machine learning methods. Post hoc analysis shows that scoring functions based only on ligand features are competitive with those also using structural features. Our DNN approach tied for fifth in predicting BACE-ligand binding affinities.

## Introduction

The Drug Design Data Resource (D3R) has organized four Grand Challenges (GC) for docking, affinity, and free energy predictions for protein-ligand complexes.[1–3] By providing high quality, blinded protein-ligand crystal structures and affinity datasets, D3RGC has attracted extensive attention from computational drug design researchers. Assessment of results from blinded competition has provided unbiased insights into the most effective strategies as well as the many shortcomings of the state of the art. This platform has sparked numerous novel computer-aided drug designing methods. In this article, we present our machine learning approach in GC4 for predicting ligand binding affinities for beta-secretase 1 (BACE), a key protein for the formation of amyloid β-peptide and a leading drug target for Alzheimer’s disease.[4]

Computer-aided drug design facilitates the whole process of drug development, such as virtual screening, lead optimization, structure-activity relationships (SAR) analysis, and absorption, distribution, metabolism, excretion, toxicity (ADMET) modeling.[5] The designing and modeling of drug molecules can be classified into structure-based drug design (SBDD) and ligand-based drug design (LBDD), depending on whether three-dimensional structural information is used.[6] To design a novel drug molecule, an accurate prediction of the binding affinity between a ligand and its target protein is helpful. However, due to the complex nature of intermolecular interactions, protein flexibility, solvation, and the entropic effect, the prediction of docked structures and affinities of protein-ligands is very challenging. Classically, a predicted docked pose needs to be generated first, then the affinity is calculated by force-field-based, knowledge-based, or an empirical scoring function. Popular docking programs and scoring functions such as the AutoDock family[7–9], Glide[10], GOLD[11], and X-Score[12] have demonstrated their advantages in predicting docked poses and protein-ligand affinities.

Machine learning models have shown great potential in advancing current computer-aided drug designing methodology in the fields of protein folding, protein binding site prediction, ligand pose and affinity prediction, and generation of drug-like molecules.[13–15] The advance of machine learning platforms and tools for chemistry, such as TensorFlow[16] and scikit-learn[17], and the public availability of high-quality protein-drug datasets, such as the PDB[18] and PDBbind[19], greatly enables the application of machine learning to drug design. Novel machine learning-based scoring methods, such as RF-Score[20] and K_DEEP_[21], or machine learning-optimized software, such as Vinardo[22], smina[23], RF-Score-v3[24], have demonstrated superior performance in addition to their generalizability and accessibility. Machine learning methods have been shown to perform better than conventional scoring methods in benchmark studies. [25] For example, machine learning scoring functions TopBP, K_DEEP_, RF-Score v.3, X-Score, and cyScore have Pearson’s correlation coefficients (R) of 0.86, 0.82, 0.80, 0.66, 0.65, respectively, for 290 protein-ligand affinities in the PDBbind v.2016 core set.[21, 26] Surprisingly, four of these scoring methods are very poor at predicting the affinities of ligands to BACE in a blind dataset, as K_DEEP_, RF-Score v.3, X-Score, and cyScore yield Pearson’s correlation coefficients (R) of −0.06, −0.14, −0.12, and 0.2, respectively, for 36 BACE ligands. It is still an unresolved question why some protein targets are more difficult than others for different algorithms and scoring functions.

The special characteristics of the different published ligand-protein affinity machine learning algorithms are their unique approaches for feature extraction and different machine learning architectures. NNScore 2.0 uses seventeen empirical-based functions for distinct binding characteristics in deep neutral networks.[27] RF-Score utilizes 36 features comprising a combination of pairwise interactions between particular protein–ligand atom types in random forests. [20] K_DEEP_ treats ligand-protein complexes as 3D voxels as inputs in six convolutional channels to catch different type of physical interactions. [21] TopBP also adopts deep convolutional neural networks, using manually constructed topological features derived from persistent homology analysis. [26] Besides ligand affinity prediction, in potential energy calculations, features of atomic environment vectors using radial and angular symmetry functions in deep neutral network have shown great promise. [28]

We are especially interested in answering the question of whether machine learning models trained on a specific target can improve the affinity prediction performance for ligands to this target. Since D3R GC4 provided a high-quality and blinded BACE affinity dataset, we took this opportunity to explore the performance of a target-trained machine learning model, using basic feature extraction methods with different model architectures. We demonstrate how the combination of structure-based and ligand-based features benefit machine learning performance. We also investigated the importance of hyper-parameter tuning and feature analysis in building and refining protein-ligand affinity models.

## Methods

### Affinity model overview

To build and test the affinity prediction performance for BACE-specific trained machine learning models, there are three essential elements: training BACE input features (X training), training BACE experimental affinities (y training), and BACE test input features (X test), **Fig. 1**. To generate input features of ligands, we compared using structure-based features and ligand-based features; thus, the method of obtaining accurate docked poses is important.

**Fig. 1.**
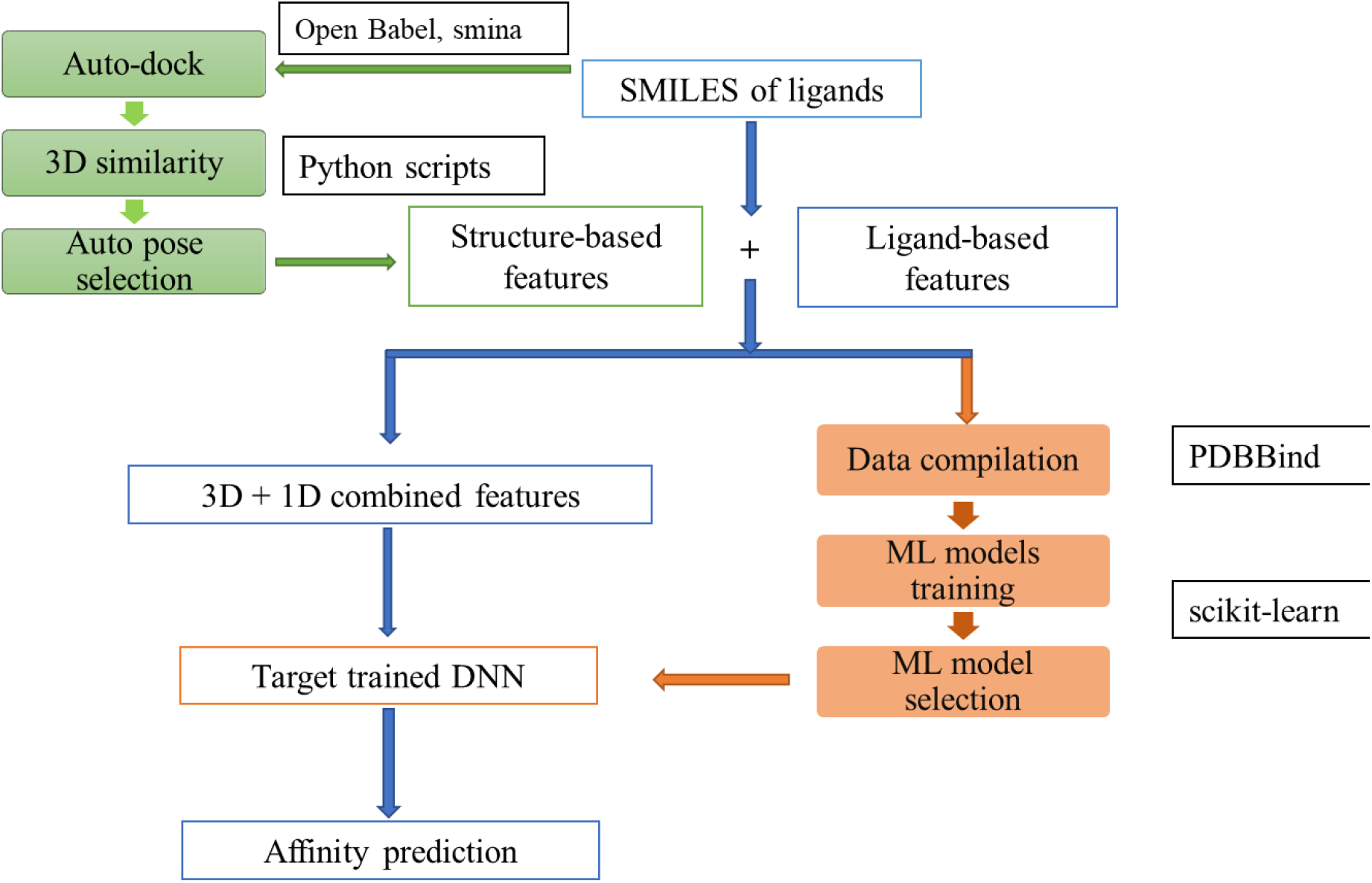
Workflow for generating machine learning models for affinity prediction. Green boxes indicate the steps in the auto-pose generator for the 154 ligands in D3R GC4 BACE Stage 2 affinity prediction challenge. Orange boxes represent procedures in the BACE-specific machine learning models of affinity prediction.

The compilation of training dataset and test input features are discussed in the “**Test dataset compilation for BACE-ligand affinity modeling**” section. Our workflow of generating predicted docked pose is introduced in the “**Semi-automated ligand pose generation and docking**” section.

### Semi-automated ligand pose generation and docking

Our first objective was to develop a docking workflow for the 154 ligands without crystal structures. It was shown in D3R GC3 that docking to the appropriate receptor structure is important for success [3]. In GC4 Stage 2, D3R provided SMILES strings of the 154 macrocyclic ligands for the affinity prediction test; 20 co-crystal structures were released earlier in D3R GC4 Stage 1B. We looked for chemical similarity between the 154 ligands (BACE_1 to BACE_158) for the affinity prediction test and the 20 ligands (BACE_1 to BACE_20) with crystal structures using the FragFp similarity method in DataWarrior 4.7.2,[29] see supplemental **Table S1** for the full similarity list. The FragFp descriptor uses chemical fingerprints based on substructure fragment matching.

We cross-docked each of the 154 test ligands to the BACE receptor bound to the ligand with the highest similarity out of BACE_1 to BACE_20. An automated cross-docking python script using Open Babel[30] and smina[23] was written to generate the docked poses for the 154 test ligands for Stage 2. Smina was used for the cross-docking procedure with the Vinardo scoring function [22]. The docking box was centered on the most similar ligand with a reduced buffer padding of 2.0 Å (default 4.0 Å) to confine docking poses in a narrower space. The reduced docking volume helped eliminate false positive poses. When multiple conformations were generated for docked ligands, large conformational differences were observed for identical ligands, indicating inadequate sampling. In such cases, we assumed that structurally similar ligand should adopt similar 3D conformations. Thus, we picked the best pose for affinity calculation based on a 3D similarity of these ligand poses to the most similar ligand structure in BACE_1 to BACE_158.

Widely used simply root-mean-square deviation (RMSD) metrics for measuring the 3D similarity for different molecules are often not applicable, since atom pairs for different compounds are not identical. To quantify the 3D similarity of multiple docked poses versus a reference ligand structure in a same receptor, we developed a 3D structural similarity score (R_sim_) inspired by crystallographic electron density difference maps. We calculated R_sim_ for each docked pose relative to the most chemical similar ligand with an available cocrystal structure:

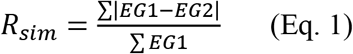

where EG denotes the electron grid of a molecule, and the summation of the electron grid in a conformation will be the total number of electrons in that molecule.

The electron grid of a molecule is calculated by directly extracting the coordinates for a docked ligand set in a box with a user-defined padding distance in each dimension (5 Å used here). The box used the same coordination system in the cocrystal. This box was then assigned with a 3D electron grid, with a grid size of 1/n Å in each dimension (n = 5 was used here, as n = 6 only has an 0.005% variation for R_sim_). To calculate the grid density, each atom of the molecule is treated as a spherical Gaussian distribution of electrons, with the integration of electron density being the atomic number, and with the standard deviation being the van der Waals radii divided by 2.3548.[31] For example, consider two poses (A and B) of the same compound. If A and B overlap with each other nearly perfectly, the electron grids of the two conformers are highly similar, thus the grids of the two conformers cancel each other, which leads to R_sim_ = 0. At the other extreme, if A and B do not spatially overlap at all (whether due to conformation, rotation, or translation), the electron grid of A has no overlap with the electron grid of B, thus the absolute value of the summation of EGA-EGB is twice the summation of EG1, thus R_sim_ = 2. An example is shown in **Fig. 2**. The Stage 2 affinity test ligand BACE_73 is most similar to BACE_10 which has a known co-crystal structure. Using smina with the Vinardo scoring function, multiple docked poses were generated for the receptor structure from the co-crystal structure of BACE_10. The second-best pose (Vinardo affinity score: −11.2 kcal/mol) and the fourth-best pose (Vinardo affinity score: −8.8 kcal/mol) are shown in **Fig. 2.** Using the R_sim_ method, the 3D-similarity of the BACE_73 second pose was shown to be more 3D-similar to BACE_10 than the BACE_73 fourth pose, as the R_sim_ is lower for the 2^nd^ pose (0.72 versus 1.36).

**Fig. 2.**
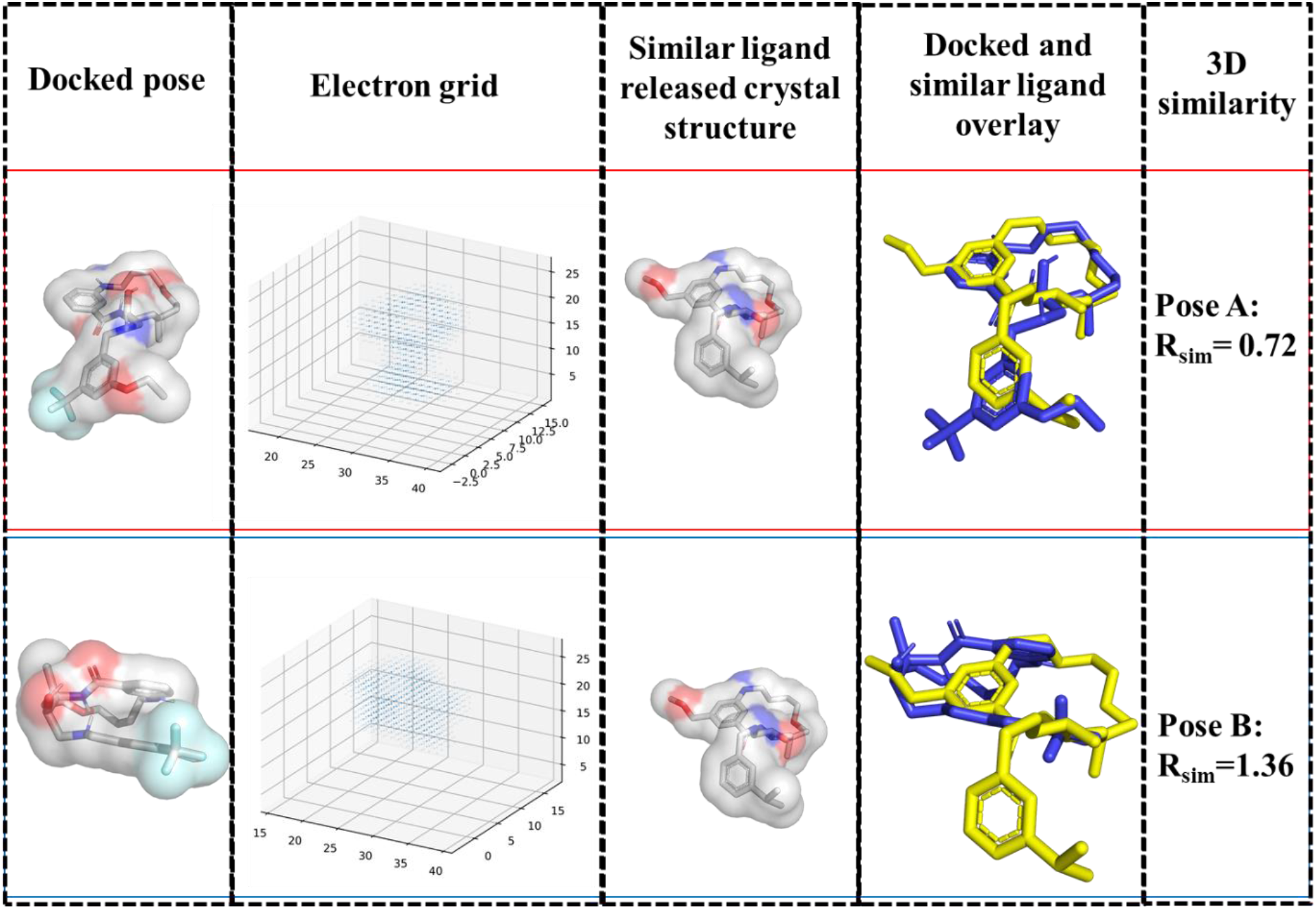
3D similarity with the R_sim_ method for D3R GC4 BACE ligands. Using the R_sim_ method, the 3D-similarity of the BACE_73 second pose (top) was shown to be more 3D-similar to BACE_10 than the BACE_73 fourth pose (bottom), as the R_sim_ is lower for the second pose. PyMOL was used to generate part of this figure. [32]

This 3D similarity algorithm worked well when comparing docked poses to cocrystal structures. For all Stage 2 ligands, the docked poses were scored with this method. Docked poses with the lowest R_sim_ value (highest 3D similarity) were selected for further analysis for affinity estimation.

### Training dataset compilation for affinity modeling of BACE-ligand

To build a BACE-specific affinity machine learning model, a dataset comprised of 222 published BACE ligand affinities (label value y_training_, dimension: 222×1) was extracted from PDBbind v.2017 which contains affinity data for 14,761 protein-ligand complexes, Table S2.[19]

In this work, structure-based and/or ligand-based features were used as input features (X) for our machine learning models. AutoDock Vina scoring functions, which combine knowledge-based potentials and empirical scoring functions, have been shown to be effective in pose prediction. [7–9] Using structural information of receptors and ligands, the Vina-like functions are derived and parameterized to estimate the strength of physical interactions, such as steric terms using Gaussian functions, and a repulsion term using a quadratic function. [23] Here we investigate whether the utilization of these already-established structure-based terms can be used as effective features for scoring the binding of a specific receptor to its ligands. To generate the structure-based features used, ten AutoDock Vina-like scoring terms were generated for the 222 BACE-ligands using smina.[23] The structure-based terms used in this work are summarized in **Table 1**. To obtain the ten scoring values, “scoring_only” functionality was used in smina, without conformation search and docking. The scores from smina were compiled as the training set (X_training_ structure-based_, dimension: 222×10).

**Table 1.** Ten structure-based AutoDock Vina terms generated by smina. Fine parameters o, w, c, g, b, s, i, j, ^ are used in the generation of the terms. Briefly, o is offset between atom pairs, w is the width of the Gaussian function, c is distance cutoff, g is a good distance, b is a bad distance, s is a smoothing parameter, i and j are Lennard-Jones exponents, ^ is the cap. [23]

| Term description | Fine parameters |
| --- | --- |
| Gaussian | o=0, w=0.5, c=8 |
| Gaussian | o=3, w=2, c=8 |
| repulsion | o=0, c=8 |
| hydrophobic | g=0.5, b=1.5, c=8 |
| van der Waals | i=6, j=12, s=1, ^=100, c=8 |
| non_dir_h_bond | g=-0.7, b=0, c=8 |
| non_dir_h_bond_lj | o= -0.7, ^=100, c=8 |
| non_dir_anti_h_bond_quadratic | o=0, c=8 |
| acceptor_acceptor_quadratic | o=0, c=8 |
| donor_donor_quadratic | o=0, c=8 |

For the ligand-based features, DataWarrior 4.7.2 was used to produce twenty-four ligand-based descriptors (X_training_ligand-based_, dimension: 222×24) for the 222 training ligands, compiled in **Table 2**. [29] When structure-based and ligand-based features were used together, a full training set (X_training_, dimension: 222×34) was obtained by combining the two datasets (X_training_structure-based_ + X_training_ligand-based_).

**Table 2.** Twenty-four ligand-based features generated by DataWarrior. [29]

| Ligand-based features | Ligand-based features |
| --- | --- |
| Molecular weight | Non-H atoms |
| cLogP | Non-C/H atoms |
| cLogS | Electronegative atoms |
| H-acceptors | Stereo-centers |
| H-donors | Rotatable bonds |
| Total surface area | Ring closures |
| Relative PSA | Small rings |
| Polar surface area | Aromatic rings |
| Drug-likeness | Aromatic atoms |
| Shape index | sp<sup>3</sup>-Atoms |
| Molecular flexibility | Symmetric atoms |
| Molecular complexity | Amides |

### Test dataset compilation for BACE-ligand affinity modeling

D3R GC4 BACE Stage 2 required the affinity prediction of 154 ligands. To utilize our BACE specifically trained machine learning model for affinity prediction, the same ten structure-based features (X_test_ structure-based_, dimension: 154 × 10) and twenty-four ligand-based features (X_test_ligand-based_, dimension: 154 × 24) for the test ligands were selected. To calculate structure-based features for the 154 test ligands, the docked poses for the ligands was generated using the method described previously. Then, the ten AutoDock Vina terms (**Table 1**) were calculated using smina. The ligand-based features for this test set were generated in the same way as the training set, using the twenty-four molecular descriptors (**Table 2**) from DataWarrior 4.7.2.

Both the X_training_ and Y_training_ were preprocessed in scikit-learn, as the data was centered to the mean and scaled to unity variance.[17] We found that when the X_training_ was not preprocessed, the machine learning models cannot be refined to convergence. When Y_training_ was not preprocessed, a similar ranking model can be obtained, however, the training process would take multiple times longer.

### Construction, training, and tuning of machine learning models for BACE-ligands affinity

After we obtained a training dataset (X_training_) and training affinities (Y_training_), five common machine learning models (**Table 3**) were constructed, refined, and compared, using structure-based and/or ligand-based features, in Python using the scikit-learn library.[17]

**Table 3.** Machine learning models for BACE-ligand affinity investigated in this study.

| Number | Model Type | Fine-tuning parameters adjusted |
| --- | --- | --- |
| 1 | Linear regression | N/A |
| 2 | Support vector machine regressor | Kernel functions, regularization (C) |
| 3 | Random forest regressor | Number of estimators, max depth |
| 4 | Regularized linear regression | Regularization ($\lambda$) |
| 5 | Deep neural network | Hidden layer size, regularization ($\lambda$) |

To validate and compare machine learning models, the training dataset was analyzed with 10-fold cross-validation with the dataset shuffled (“random state” defined as one, for reproducibility). To refine each model, hyperparameter fine-tuning of machine learning models was scrutinized and adjusted. The tuning parameters are shown in **Table 3**. For evaluation, the coefficients of determination (R^2^) with 10-fold cross-validation for each model was compared.

## Result and Discussion

### Pose generator and docking performance evaluation

In the D3R GC4 BACE Subchallenge, we also submitted pose predictions for Stages 1A (cross-docking for BACE_1 through BACE_20, with no receptor coordinates of the 20 ligands provided) and Stage 1B (redocking for BACE_1 through BACE_20, with receptor coordinates of the 20 ligands provided). We used a strategy based on optimizing the AutoDock Vina algorithm and scoring method which has been shown to perform well for pose ranking[9], where docked poses are scored based on a linear combination of five terms:

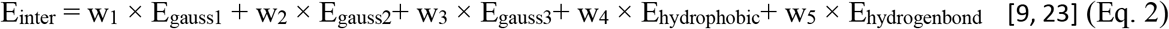

To achieve an improved cross-docking pose prediction performance for BACE specifically, the weights (wn) in the Vina scoring function (**Equation 2**) were optimized and customized on a training dataset of 24 BACE ligands and test dataset of 229 BACE ligands deposited in the PDB. The weights of the five Vina terms were refined via partial gradient descent of each weight until the overall RMSD reached a local minimum (**Table 4**).

**Table 4.** Cross-docking performance evaluation for BACE-ligands. The weights (w_n_) are the scaling factors in Equation (2). The mean and standard error of RMSD for each method were evaluated on 229 BACE-ligand cocrystal structures deposited in the PDB using smina (receptor PDB used: 4L7G).

| | $w_1$ | $w_2$ | $w_3$ | $w_4$ | $w_5$ | Mean(SE) of<br>RMSD (Å) |
| --- | --- | --- | --- | --- | --- | --- |
| **Vina[9]** | -0.035579 | -0.005156 | 0.840245 | -0.035069 | -0.587439 | 2.42(10) |
| **Vinardo<sup>a</sup>[22]** | -0.045 | 0.000 | 0.800 | -0.035 | -0.600 | 2.55(10) |
| **BACE**<br>**custom** | -0.02558 | -0.00516 | 1.19025 | -0.05507 | -0.83744 | 2.33(10) |

<sup>a</sup>Vinardo also has modified term functions besides weight.[22]

Although a statistically improved cross-docking performance was achieved for 229 BACE ligands deposited in PDB, we discovered that the Vinardo scoring function yielded the best redocking performance (lowest RMSD) for the macrocyclic BACE ligands that were the subject of D3R GC4 (**Table 5**). Thus, Vinardo scoring was used for D3R GC4 BACE Stage 1B for BACE_1 to BACE_20 redocking. The redocking performance of our method was released by D3R. Our strategy had a mean (standard deviation) RMSD of 1.97 (1.55) Å for the 20 BACE macrocyclic ligands, ranking 48^th^ place among 70 submissions. A RMSD below 2.0 Å has traditionally been considered a good result for docking. However, other research teams demonstrated superior cross-docking and redocking performance during this challenge. We expect that more thorough conformer sampling and an additional relaxation and optimization step after docking would improve our docking performance. The excellent docking performance of D3R participants suggest that at least for BACE, ligand docking is not nearly as difficult as affinity prediction.

**Table 5.**
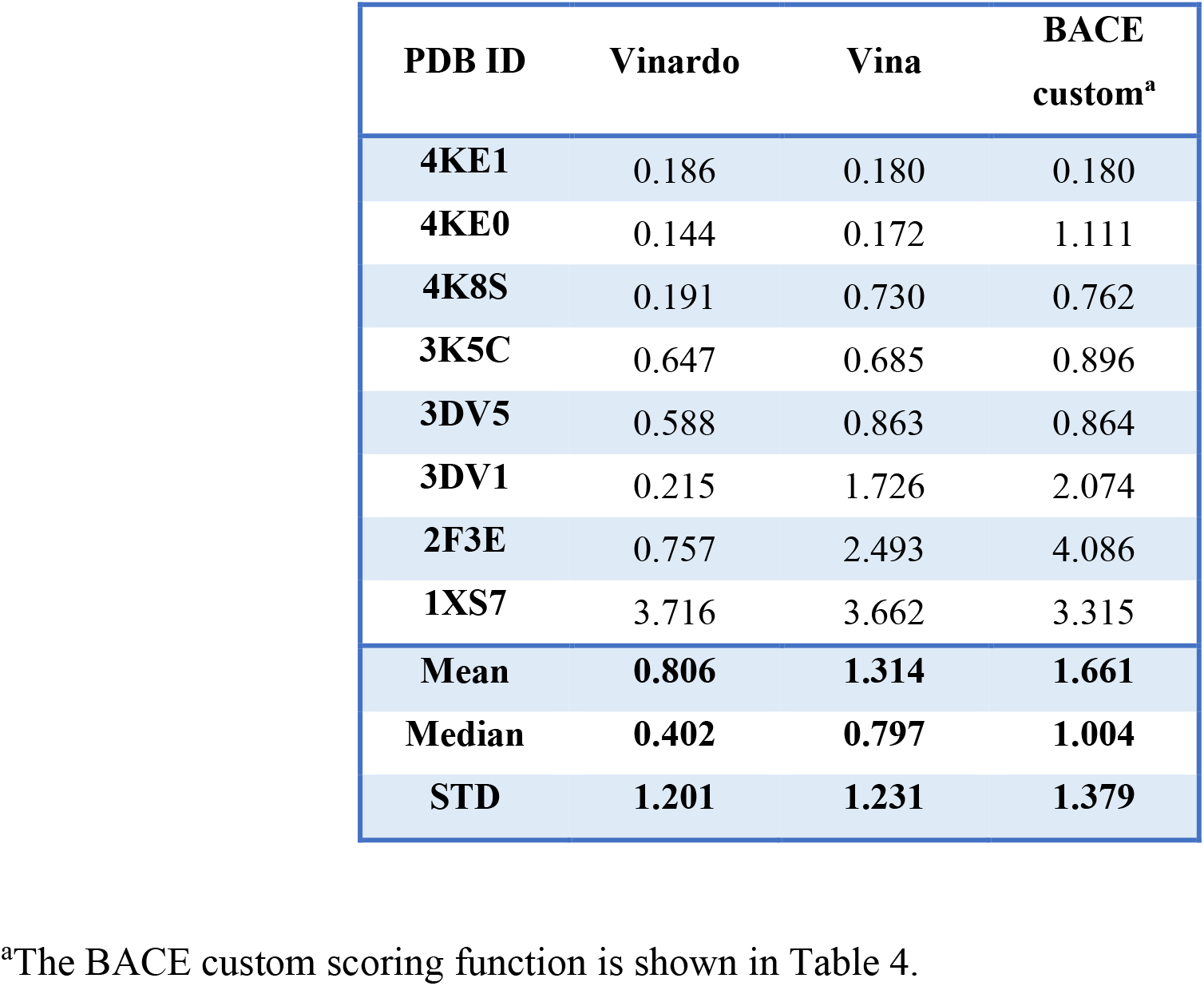
Redocking performance evaluation for BACE-macrocyclic ligands deposited in PDB. Docking was conducted with smina with Vina/Vinardo/custom scoring methods. [22] Test cocrystal structures were obtained from the PDB. RMSD was calculated and compared using PyMOL. Vinardo has the lowest mean RMSD, indicating the best pose ranking performance.

| **PDB ID** | **Vinardo** | **Vina** | **BACE custom<sup>a</sup>** |
| --- | --- | --- | --- |
| **4KE1** | 0.186 | 0.180 | 0.180 |
| **4KE0** | 0.144 | 0.172 | 1.111 |
| **4K8S** | 0.191 | 0.730 | 0.762 |
| **3K5C** | 0.647 | 0.685 | 0.896 |
| **3DV5** | 0.588 | 0.863 | 0.864 |
| **3DV1** | 0.215 | 1.726 | 2.074 |
| **2F3E** | 0.757 | 2.493 | 4.086 |
| **1XS7** | 3.716 | 3.662 | 3.315 |
| **Mean** | **0.806** | **1.314** | **1.661** |
| **Median** | **0.402** | **0.797** | **1.004** |
| **STD** | **1.201** | **1.231** | **1.379** |

<sup>a</sup>The BACE custom scoring function is shown in Table 4.

After D3R GC4 Stage1B, twenty cocrystal structures of BACE_01:BACE_20 macrocyclic ligands were released. When overlaying the D3R ligands with macrocyclic ligands deposited in PDB, it was found that these ligands shared a similar structural motif in the main site: the macrocycle occupying an empty cavity with a large substituent extended to the other side of the main cavity via a linker that usually contains hydrogen-binding, electronegative functional groups, as shown in **Fig 3**.

**Fig. 3.**
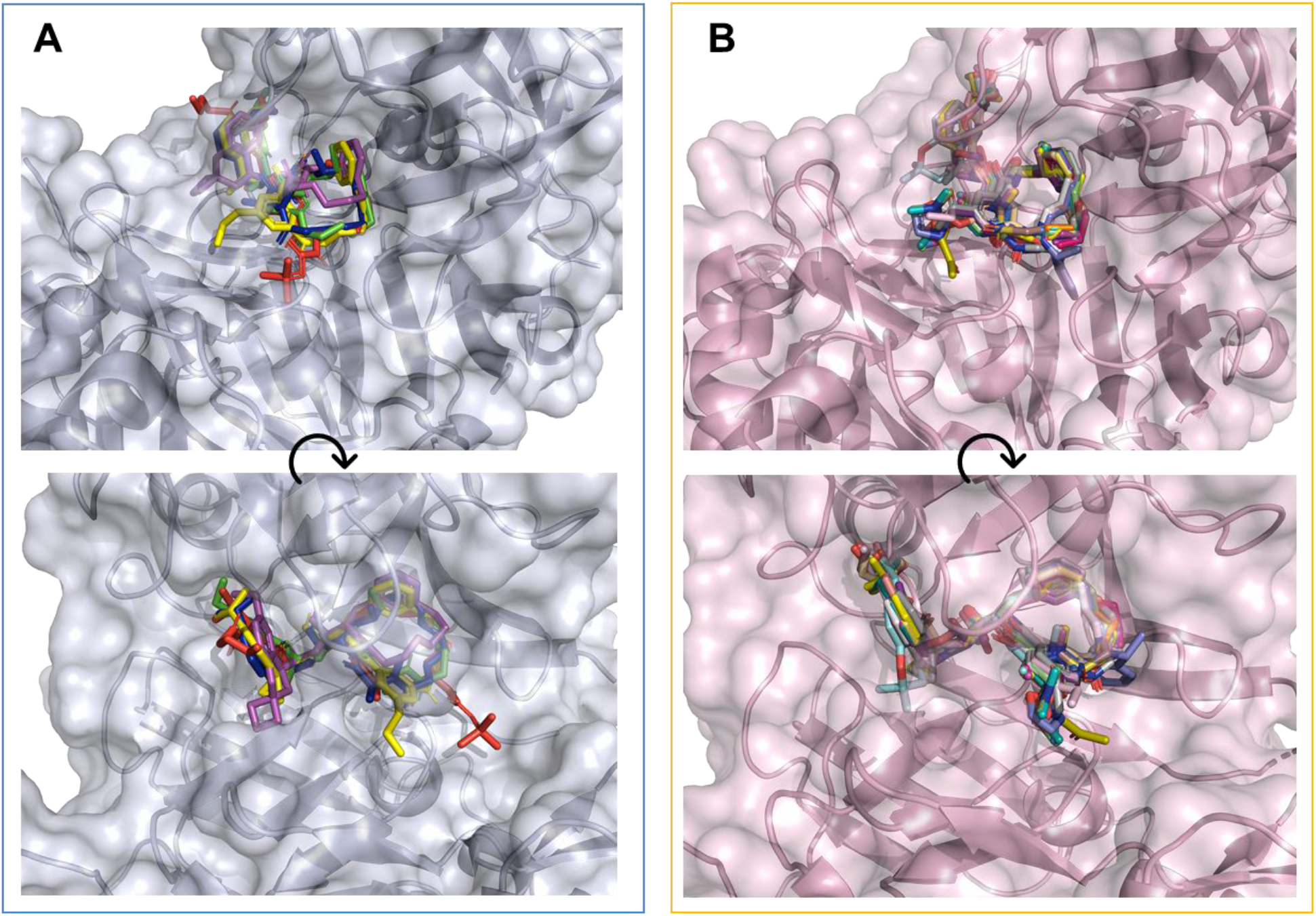
Structure overlay of BACE macrocyclic ligands. A, five published ligands structures deposited in the PDB: 2F3E-AXQ(red), 3DV1-AR9(green), 3DV5-BAV(blue), 3K5C-OBI(yellow), 4KE0-1R8(magenta), receptor PDB used: 4KE0.[18] B, twenty D3R GC4 released macrocyclic ligands (BACE_1 to BACE_20, receptor used BACE_BA01. Top and bottom figures show rotated views of the same structure. PyMOL was used to generate this figure.

Since the Vinardo scoring function was shown to be effective in generating docked poses for macrocyclic BACE ligands (**Table 5**), we used an automated pose selection method to select the best pose from the multiple docked poses generated by smina/Vinardo.

We hypothesized that chemically similar BACE macrocyclic ligands should share similar docked poses (154 BACE chemical similarity pairs are provided in **Table S1**). The semi-automated docking and pose selection workflow was described above. Given multiple cross-docked poses of the 154 ligands without crystal structures, the pose with the lowest R_sim_ (defined in **Equation 1**) was selected for structure-based scoring. Two examples are shown in **Fig 4**. BACE_26 (test) is chemically similar to BACE_3 (crystal structure known). A docked pose generated by smina/Vinardo produced a lower R_sim_ = 0.553 and was used for affinity prediction. BACE_137 is similar to BACE_12. A docked pose with R_sim_ = 0.564 was used for further calculation. Using this semi-automated workflow, 154 predicted docked structures were generated for BACE-ligand affinity prediction. We expect that further optimization of the selected pose will improve pose accuracy.

**Fig. 4.**
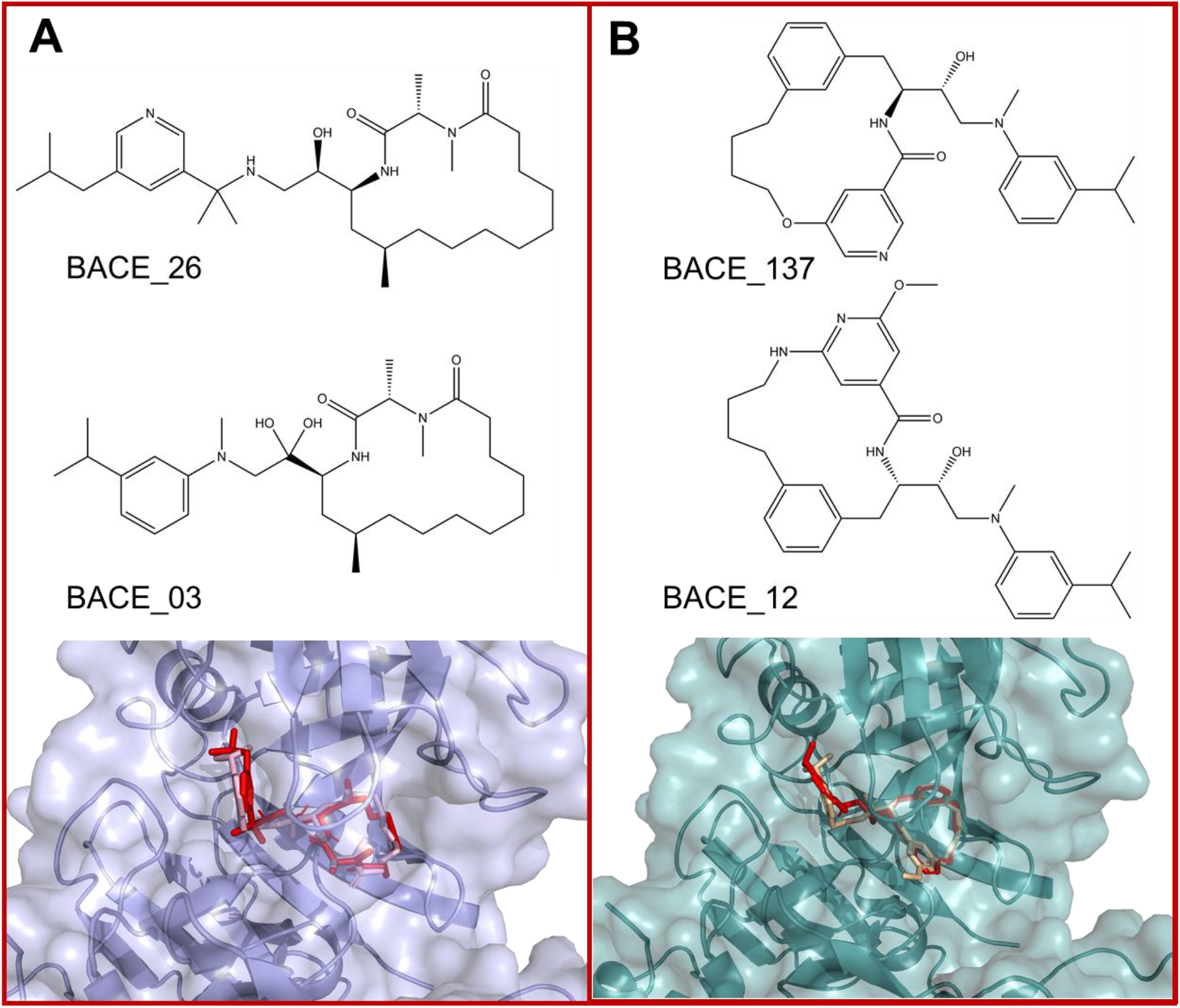
Chemical structures and structural overlays of two sets of docked macrocyclic ligands in D3R GC4 BACE Stage 2. A, BACE_26 (red) was found to be most chemically similar to BACE_3 (light pink), that has a released cocrystal structure, via the FragFp method in DataWarrior. Between two docked poses of BACE_26 generated by Vinardo scoring using smina, the pose with the lowest R_sim_ = 0.553 was used for structure-based affinity calculations. B, similarly, ligand BACE_137 (red) was docked to mimic the 3D structure of the most chemically similar ligand BACE_12 (light orange); the pose with the lowest R_sim_ = 0.564 was used for affinity modeling. ChemBioDraw and PyMOL were used to generate this figure.

### Comparisons between machine learning models of BACE-ligand affinities

We obtained a dataset (X_training_) of 222 BACE-ligands deposited in PDB with their affinities (Y_training_) extracted from PBDBind v2017[19]. We investigated three aspects of applying machine learning in BACE-ligand affinity prediction: input feature selection, machine learning model selection, and machine learning model regularization.

First, for feature selection, we compared model performance based on structure-based features only **(Table 1**), ligand-based features only (**Table 2**), and a combination of both using five machine learning models (**Table 3**): linear regression, regularized linear regression, random forest regressor (RF), support vector machine regressor (SVM), and deep neural network/multilayer perceptron (DNN/MLP).

When only structure-based terms were used for affinity modeling (**Fig. 5A**), linear regression (LR) and regularized LR (λ = 10) yielded poor performance, with low coefficients of determination (R^2^) at 0.30(7), 0.28(8) for the ten-fold cross-validation set (X_training_). This indicates the limitations of linear customization of Vina terms to predict ligand affinity. We point out that multilinear regression is still the most commonly used method for developing docking and affinity scoring functions. Using random forest regressor (RF, number of estimators: 200, maximum depth: 8), support vector machine regressor (SVM, kernel: Gaussian radial basis function, regularization: C = 2), and multiple layer perceptron (MLP, layer structure: 10 × 8 × 8, activation function: ReLU, L2 regularization: alpha = 10) with structure-based only terms yielded reasonable performance after fine-tuning the hyperparameters. The SVM regressor model performed well and achieved an R^2^ of 0.65(11) for the 10-fold cross-validation set, and 0.77(1) for the same 10-fold cross-training set. Our results strongly support the use of modern non-linear machine learning methods scoring functions for predicting ligand affinity.

**Fig. 5.**
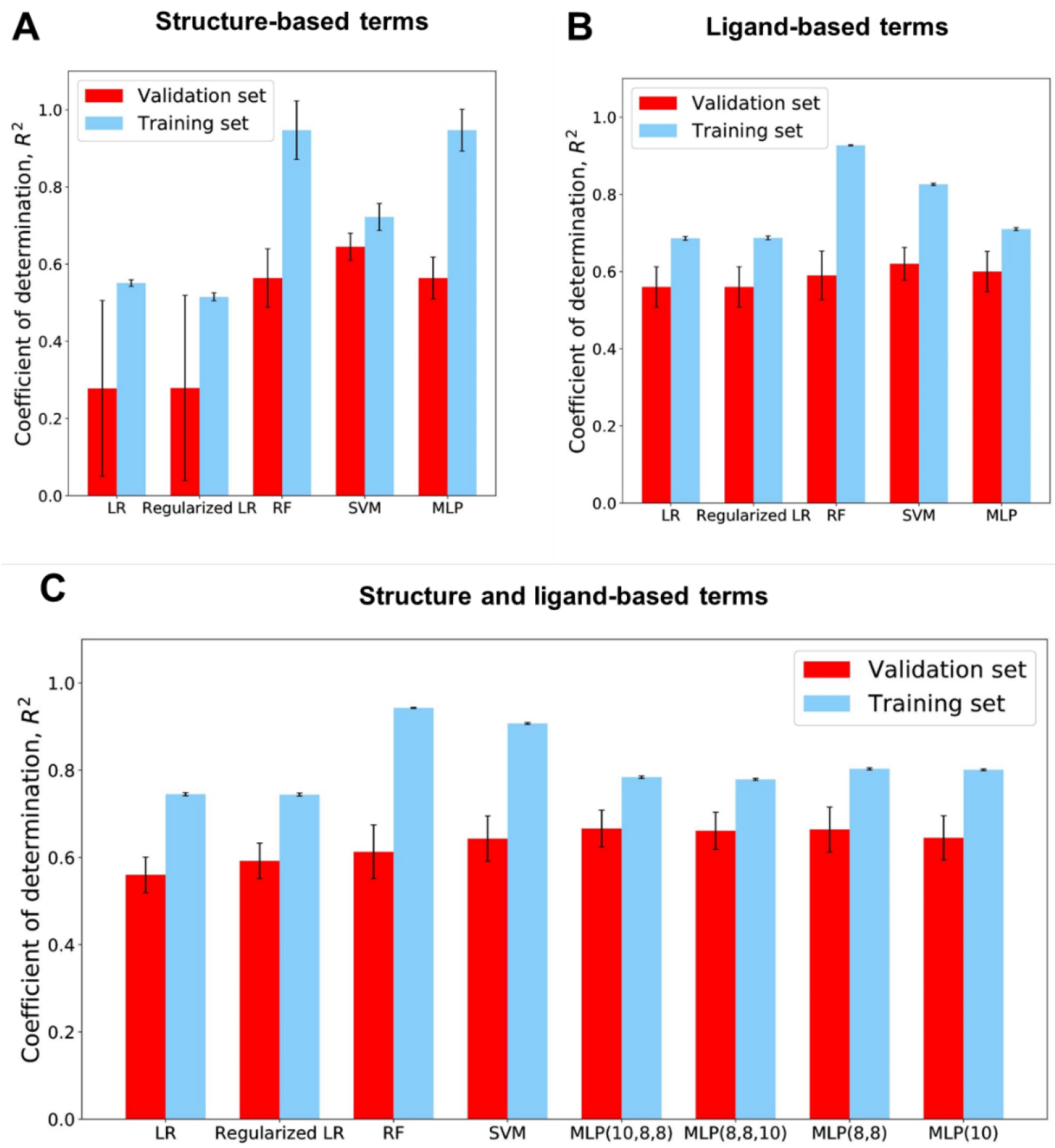
Performance of machine learning models of BACE affinity. Red boxes indicate the mean of R^2^ for the 10-fold cross-validations, cyan boxed indicate the mean of R^2^ for the training set. The error bars represent the standard error (SE) for the 10-fold cross-validation metric evaluation. A, only the ten structure-based Vina terms were used. B, only the twenty-four ligand-based terms (Table 2) were used. C, both the structure-based and ligand-based terms were applied. Numpy, Matplotlib, and scikit-learn were used to generate this figure.

The performance of only using ligand-based terms was also evaluated (**Fig. 5B)**. Interestingly, all five machine learning models yielded comparable results, suggesting that linear regression adequately utilizes most of the information in the input features. The SVM performed the best among the models, with an R^2^ of 0.62(13) for the 10-fold cross-validation set and 0.836(9) for the cross-training set.

When a combination of structure-based terms and ligand-based terms was utilized in machine learning models, slightly improved performance was obtained across the different modeling methods (**Fig. 5C**). LR and regularized LR (λ = 10) yielded equivalent performance, with R^2^ = 0.59(13) for the 10-fold cross-validation set. RF (number of estimators: 200, maximum depth: 8) yielded an R^2^ of 0.61(19) for the cross-validation set. SVM regressor (kernel: Gaussian, regularization: C = 2.5) had a R^2^ = 0.64(17) for the validation set. We compared DNN/MLP architectures with 10 × 8 × 8, 8 × 10 × 10, 8 × 8, and 10 (**Table 6)**. Only four networks were explored given limits on our time. It was found that the MLP model performance is slightly affected by the choice of the number of layers and numbers of neurons in layers, as R^2^ of 0.65 were achieved in all four architectures. Careful hyperfine tuning is necessary to obtain an effective model. For example, in the 10 × 8 × 8 DNN, an alpha (L2 regularization term) of 0.01 overfit the training set and yielded a poor R^2^ (0.3±0.4) of the ten-fold cross-validation sets; when alpha of 50 was used, the model was underfitted, with poor R^2^ (0.54±0.15).

For the D3R GC4 BACE Stage 2 affinity predictions, we selected the refined 10 × 8 × 8 DNN model due to its superior performance over other architectures. The hyperparameter tuning of this model is shown in **Fig. 6A**. With the L2 regularization parameter alpha = 10, the model achieves optimal performance on the ten-fold cross-validation, with Pearson’s correlation (R) = 0.82 (R^2^ = 0.67). Using this model, the predicted versus experimental affinities for the whole training dataset are shown in **Fig. 6B** (the last cross-validated model in the 10-fold cross-validation dataset). This model exhibited very good correlation metrics for BACE affinity prediction. It greatly outperforms the published performance of K_DEEP_, RF-Score, X-Score, and Cyscore, with their R equal to −0.06, −0.14, −0.12, and 0.2, respectively to 36 BACE ligands[21]. The architecture and mapping matrix (Wn) are represented in **Fig. 6C**. Every neuron in the MLP take a linear combination of earlier neurons as input, and output after ReLU (rectified linear unit) activation.

**Fig. 6.**
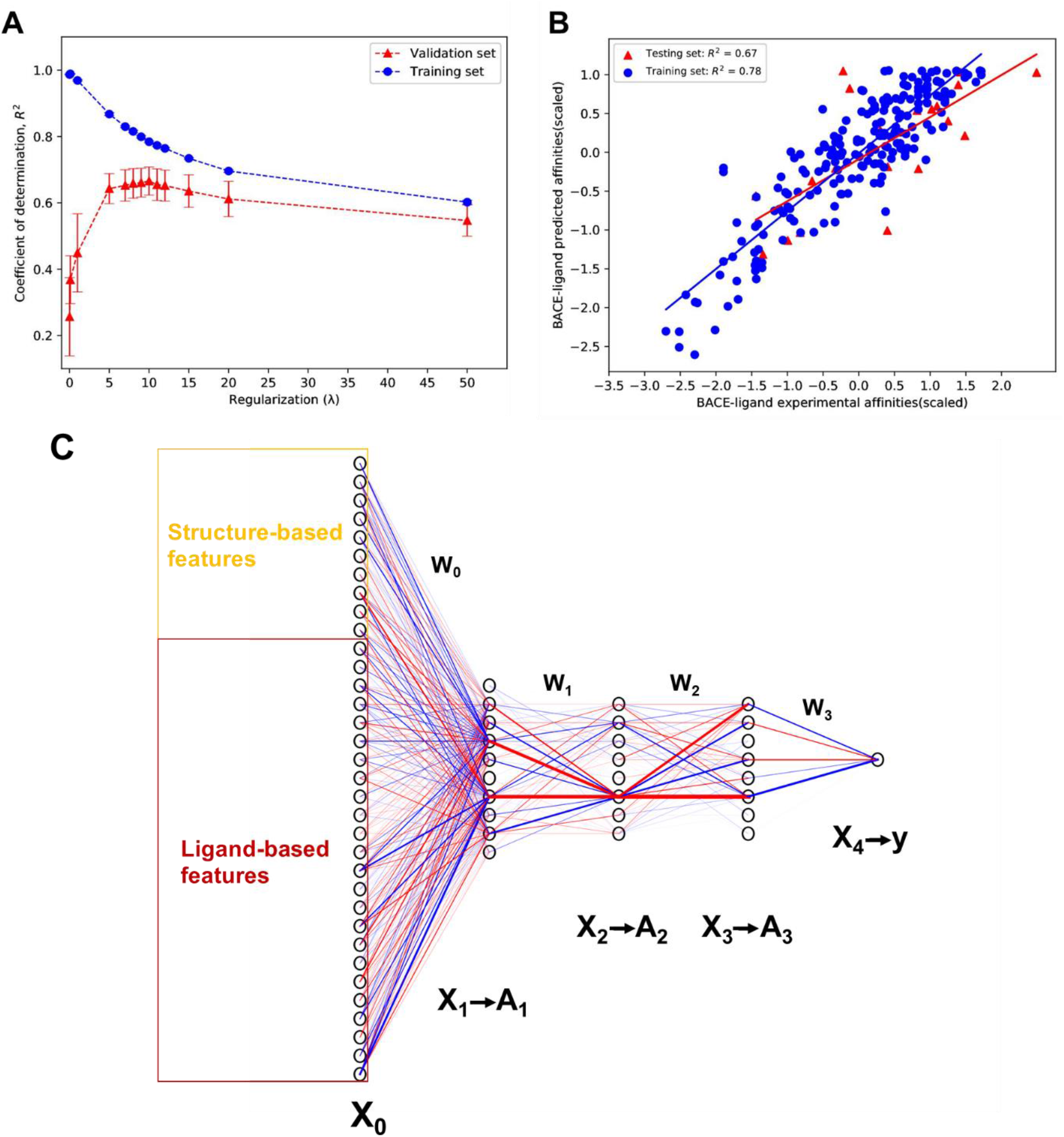
The regularization, fitting, and representation of our DNN/MLP model submitted for D3R GC4 BACE Stage 2. Model parameters: layer structure: 10 × 8 × 8, activation function: ReLU, regularization: alpha = 10. A, MLP model performance (R^2^) over variations of a range of L2 regularization alpha (0.01-50.0). B, BACE affinity fitting ability of this MLP model. Blue dots are for the last training set in the 10-fold cross-training, and red spots are for the last validation set in 10-fold cross-validation. C, the architect and mapping coefficient of this MLP. Redlines indicate positive coefficients and blue lines indicate negative coefficients, and the thickness indicates the absolute value of coefficients connecting neurons. Numpy, Matplotlib, and scikit-learn were used to generate this figure.

D3R released the performance of GC4 BACE Stage 2. Our results tied for the fifth best performance for the BACE affinity prediction in combined ligand- and structure-based scoring. The submitted MLP model has Kendall’s τ = 0.30(5), Spearman’s p = 0.43(7), and Pearson’s r = 0.45. To compare within our models, they were evaluated against the D3R GC4 BACE Stage 2 dataset. The performance and parameters of each model are summarized in **Table 7** and visualized in **Fig. 7**. Against the D3R GC4 BACE test set, the ligand-based models perform better than the structure-based models. The combination of structure and ligand-based models do not offer apparent enhancement of model performance. In fact, both the ligand-based SVM and ligand-based MLP model might slightly outperform the MLP model submitted, as the two ligand-based models has Kendall’s τ = 0.33(5) and Spearman’s p = 0.47(7). This evaluation suggests that the incorporation of structure-based features do not enhance the accuracy of affinity prediction of BACE - macrocycle ligands. Feature analysis described in the section below further demonstrates the lack of benefit from incorporating those structure-based features in predicting the affinity of macrocyclic ligands of BACE.

**Fig. 7.**
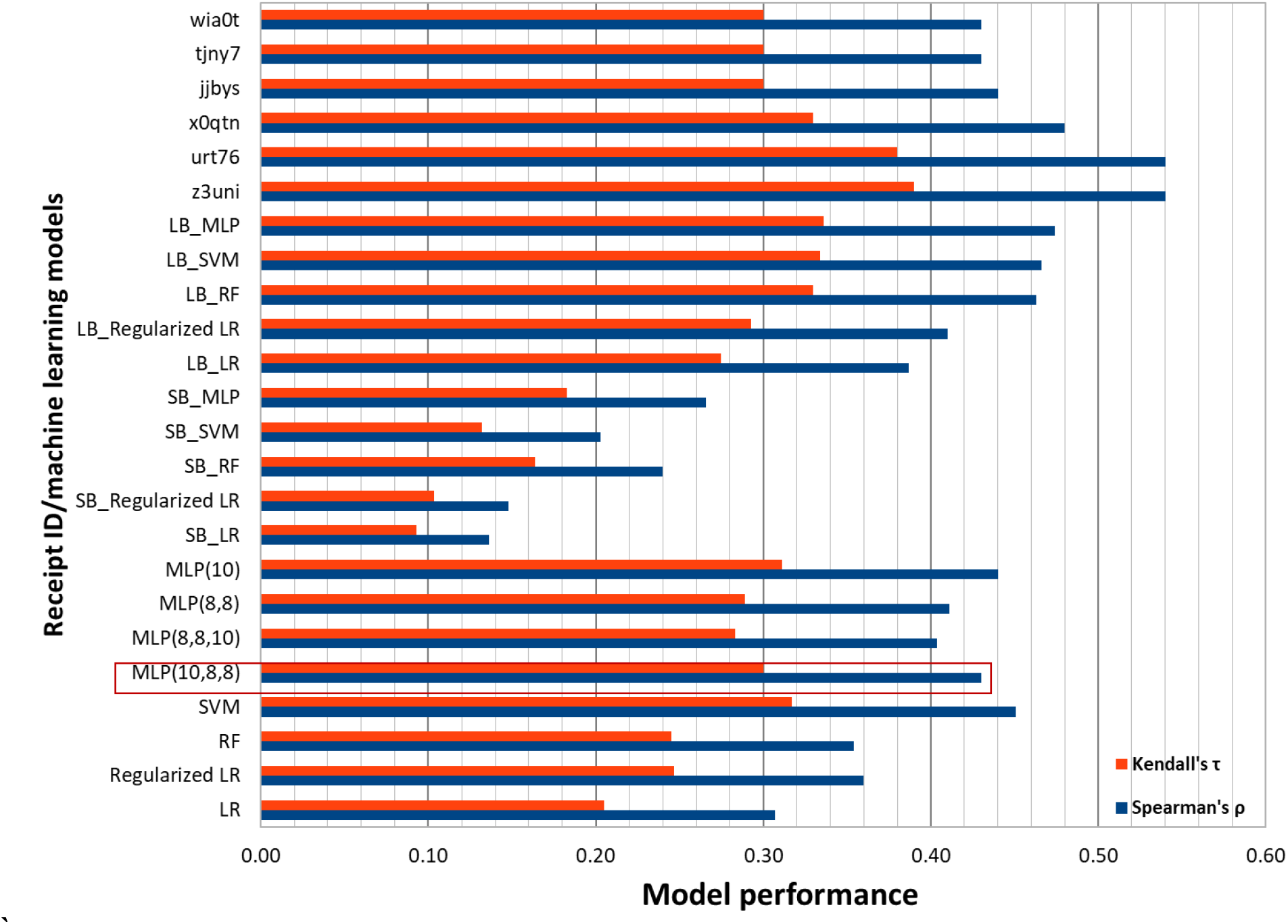
Model performance comparison against the GC4 BACE Stage 2 dataset, using Spearman’s ρ (blue bars) and Kendall’s τ (red bars). The top 6 models released by D3R are placed at the top, as indicated by their receipt ID. All the machine learning models built in this work were evaluated against the GC4 BACE Stage 2 models after the results were released. Ligand-based methods have model names starting with “LB”, and structure-based methods have model names starting with “SB”. The eight models at the bottom are the ligand- and structure-based model. Our MLP(10,8,8) model submitted to D3R is indicated by the red rectangle.

**Table 6.**
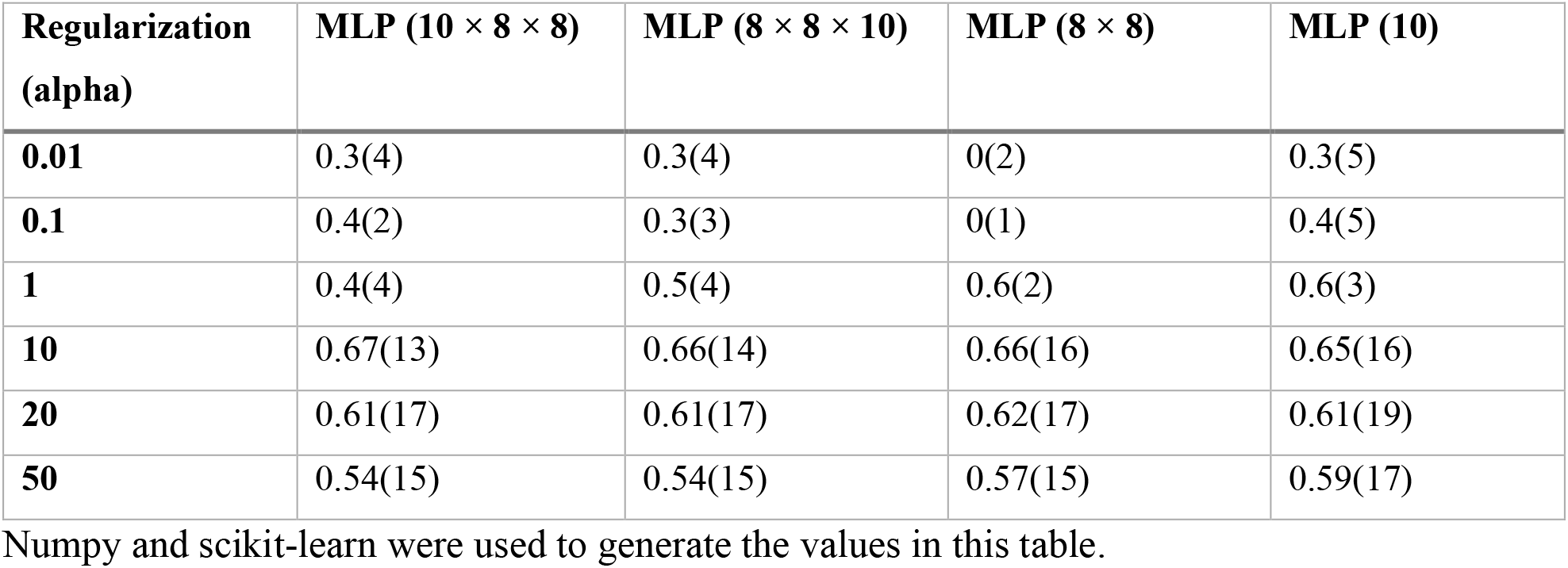
Evaluation (R^2^ of the validation sets) of different layer structures of MLP models against 10-fold cross-validation for the 222 BACE-ligands dataset (X_training_). Activation function: ReLU, L2 regularization parameter was adjusted. Mean of the R^2^ is shown here, and the number in the bracket is the standard derivation for the 10-fold cross-validation.

**Table 7.**
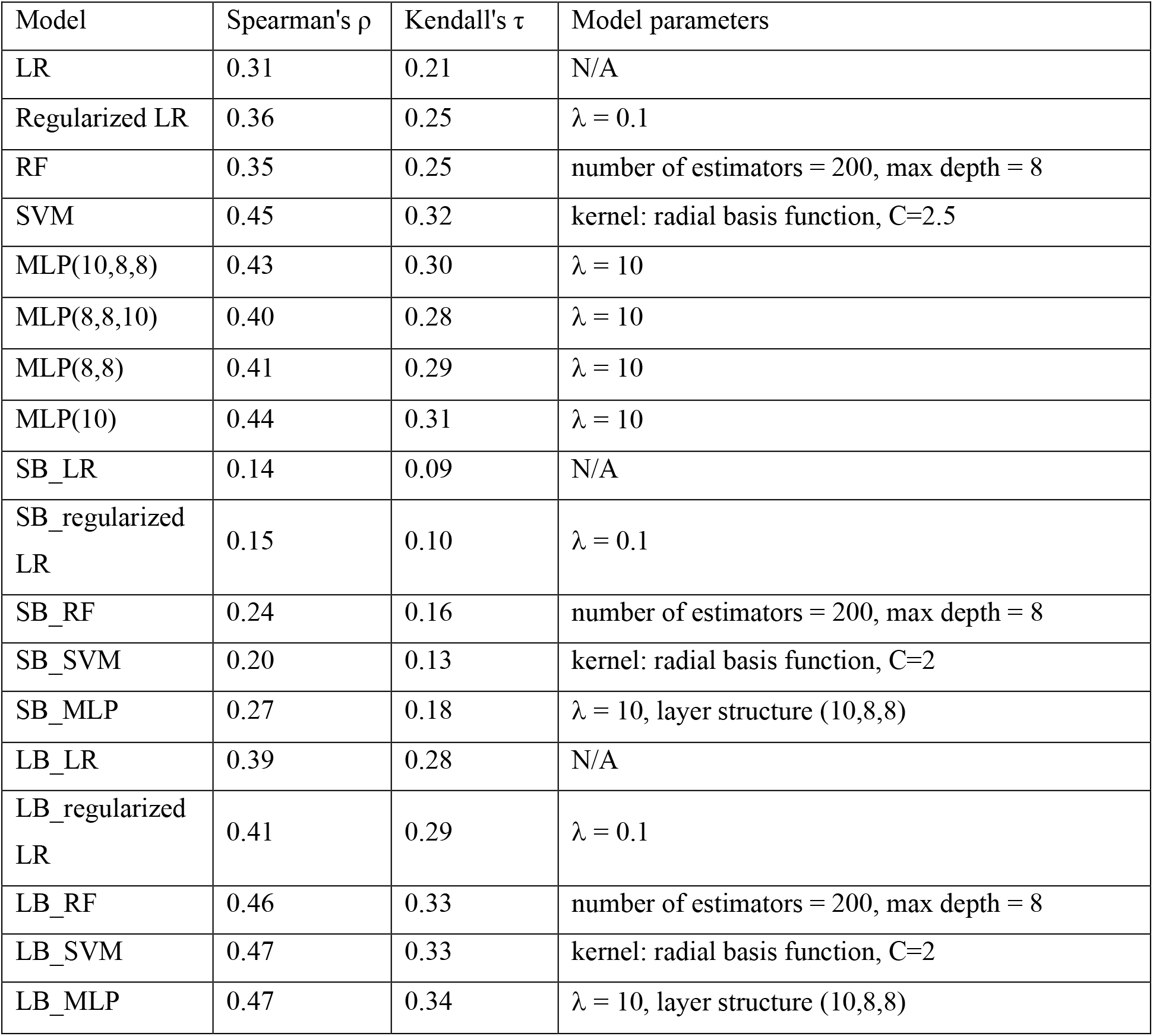
Performance and model parameters of the proposed machine models. Models were evaluated against the D3R GC4 BACE test set (sample size = 154). Model names starting with “SB” are for structure-based models, and “LB” are for ligand-based model. Model names without “SB” or “LB” are the combined structure- and ligand-based models. The MLP(10,8,8) model is the one we submitted to D3R GC4 BACE stage 2.

| Model | Spearman's $\rho$ | Kendall's $\tau$ | Model parameters |
| --- | --- | --- | --- |
| LR | 0.31 | 0.21 | N/A |
| Regularized LR | 0.36 | 0.25 | $\lambda = 0.1$ |
| RF | 0.35 | 0.25 | number of estimators = 200, max depth = 8 |
| SVM | 0.45 | 0.32 | kernel: radial basis function, C=2.5 |
| MLP(10,8,8) | 0.43 | 0.30 | $\lambda = 10$ |
| MLP(8,8,10) | 0.40 | 0.28 | $\lambda = 10$ |
| MLP(8,8) | 0.41 | 0.29 | $\lambda = 10$ |
| MLP(10) | 0.44 | 0.31 | $\lambda = 10$ |
| SB_LR | 0.14 | 0.09 | N/A |
| SB_regularized LR | 0.15 | 0.10 | $\lambda = 0.1$ |
| SB_RF | 0.24 | 0.16 | number of estimators = 200, max depth = 8 |
| SB_SVM | 0.20 | 0.13 | kernel: radial basis function, C=2 |
| SB_MLP | 0.27 | 0.18 | $\lambda = 10$, layer structure (10,8,8) |
| LB_LR | 0.39 | 0.28 | N/A |
| LB_regularized LR | 0.41 | 0.29 | $\lambda = 0.1$ |
| LB_RF | 0.46 | 0.33 | number of estimators = 200, max depth = 8 |
| LB_SVM | 0.47 | 0.33 | kernel: radial basis function, C=2 |
| LB_MLP | 0.47 | 0.34 | $\lambda = 10$, layer structure (10,8,8) |

### Feature analysis

After the D3R BACE result was released, we conducted feature analysis to gain a better understanding of the machine learning models we trained for BACE-ligand affinity prediction.

We analyzed the correlation between the features utilized and the affinity of the training set (sample size: 222), and to the D3R BACE test set (sample size: 154). The Pearson’s r and Spearman’s ρ for the feature-affinity correlation are compiled in **Table 8**. In the BACE training set, six out of ten structure-based features show moderate to strong positive or negative correlation (ρ > 0.40 or < −0.40) to the affinity. However, in the D3R BACE test set, none of the structure-based features shows moderate correlation to the affinity, with the highest ρ = 0.39 for feature Gaussian2. For ligand-based features, eleven out of twenty-four features show moderate to strong positive or negative correlation in the training set. Only four ligand-based features show moderate correlation in the D3R BACE test set, namely molecular weight (ρ = 0.43), total surface area (ρ = 0.45), number of non-H atoms (ρ = 0.40), and symmetric atoms (ρ = 0.42). This reduced correlation in the D3R test set suggests a weakened differentiation ability of macrocyclic ligands using these ten structure-based and twenty-four ligand-based features. Indeed, in the training set, where the BACE ligands are chemically diverse, those basic features are very effective to relate to their affinity. However, in the D3R test set, where the ligands are all macrocyclic and relatively similar, the features we used are less effective to differentiate those test ligands. Furthermore, in the training set, the cocrystal structures are known. But the ligands’ conformations in the D3R test set were derived in silico. Thus, the structure-based feature in the test set has underlying uncertainties and errors that are likely to reduce the feature-affinity correlation. Note that the four features alone (molecular weight, total surface area, number of non-H atoms, and symmetric atoms) can rank the BACE-ligand affinity moderately well. For example, the total surface energy and affinity is strongly related in the training set (Spearman’s ρ = 0.62), and already slightly outperform our MLP(10,8,8) model in the D3R test set (Spearman’s ρ = 0.45).

**Table 8.**
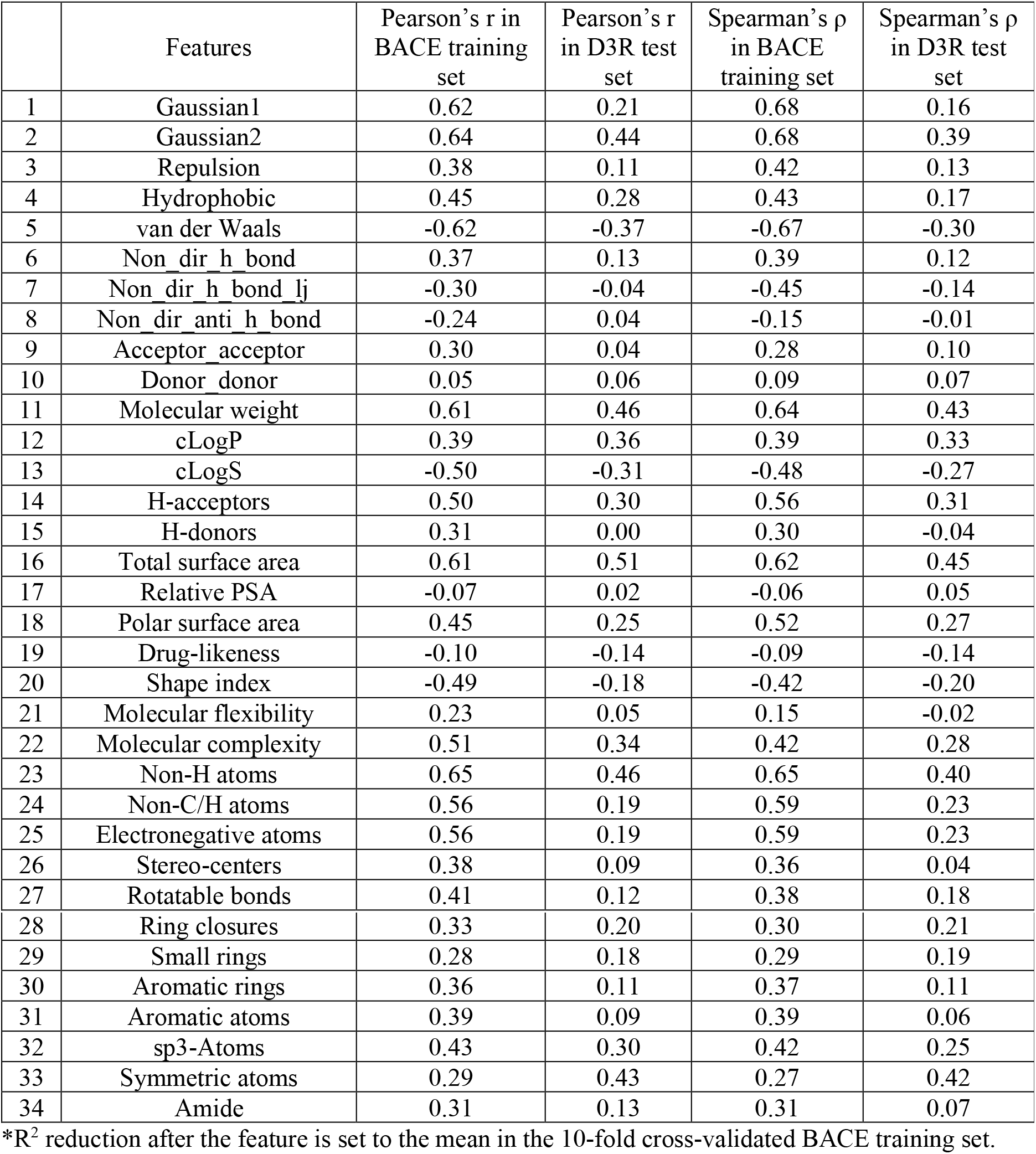
Normalized feature-affinity correlation in BACE training set (sample size = 222) and D3R GC4 BACE stage 2 (sample size = 154).

| | Features | Pearson's r in BACE training set | Pearson's r in D3R test set | Spearman's $\rho$ in BACE training set | Spearman's $\rho$ in D3R test set |
| --- | --- | --- | --- | --- | --- |
| 1 | Gaussian1 | 0.62 | 0.21 | 0.68 | 0.16 |
| 2 | Gaussian2 | 0.64 | 0.44 | 0.68 | 0.39 |
| 3 | Repulsion | 0.38 | 0.11 | 0.42 | 0.13 |
| 4 | Hydrophobic | 0.45 | 0.28 | 0.43 | 0.17 |
| 5 | van der Waals | -0.62 | -0.37 | -0.67 | -0.30 |
| 6 | Non_dir_h_bond | 0.37 | 0.13 | 0.39 | 0.12 |
| 7 | Non_dir_h_bond_lj | -0.30 | -0.04 | -0.45 | -0.14 |
| 8 | Non_dir_anti_h_bond | -0.24 | 0.04 | -0.15 | -0.01 |
| 9 | Acceptor_acceptor | 0.30 | 0.04 | 0.28 | 0.10 |
| 10 | Donor_donor | 0.05 | 0.06 | 0.09 | 0.07 |
| 11 | Molecular weight | 0.61 | 0.46 | 0.64 | 0.43 |
| 12 | cLogP | 0.39 | 0.36 | 0.39 | 0.33 |
| 13 | cLogS | -0.50 | -0.31 | -0.48 | -0.27 |
| 14 | H-acceptors | 0.50 | 0.30 | 0.56 | 0.31 |
| 15 | H-donors | 0.31 | 0.00 | 0.30 | -0.04 |
| 16 | Total surface area | 0.61 | 0.51 | 0.62 | 0.45 |
| 17 | Relative PSA | -0.07 | 0.02 | -0.06 | 0.05 |
| 18 | Polar surface area | 0.45 | 0.25 | 0.52 | 0.27 |
| 19 | Drug-likeness | -0.10 | -0.14 | -0.09 | -0.14 |
| 20 | Shape index | -0.49 | -0.18 | -0.42 | -0.20 |
| 21 | Molecular flexibility | 0.23 | 0.05 | 0.15 | -0.02 |
| 22 | Molecular complexity | 0.51 | 0.34 | 0.42 | 0.28 |
| 23 | Non-H atoms | 0.65 | 0.46 | 0.65 | 0.40 |
| 24 | Non-C/H atoms | 0.56 | 0.19 | 0.59 | 0.23 |
| 25 | Electronegative atoms | 0.56 | 0.19 | 0.59 | 0.23 |
| 26 | Stereo-centers | 0.38 | 0.09 | 0.36 | 0.04 |
| 27 | Rotatable bonds | 0.41 | 0.12 | 0.38 | 0.18 |
| 28 | Ring closures | 0.33 | 0.20 | 0.30 | 0.21 |
| 29 | Small rings | 0.28 | 0.18 | 0.29 | 0.19 |
| 30 | Aromatic rings | 0.36 | 0.11 | 0.37 | 0.11 |
| 31 | Aromatic atoms | 0.39 | 0.09 | 0.39 | 0.06 |
| 32 | sp3-Atoms | 0.43 | 0.30 | 0.42 | 0.25 |
| 33 | Symmetric atoms | 0.29 | 0.43 | 0.27 | 0.42 |
| 34 | Amide | 0.31 | 0.13 | 0.31 | 0.07 |

\*R<sup>2</sup> reduction after the feature is set to the mean in the 10-fold cross-validated BACE training set.

Besides feature-affinity correlation, the normalized covariance matrices of features in the training set and the D3R test set were generated and shown in **Fig. 8**. In the covariance matrices, the value of each element indicates the degree and direction of the relationship between two feature variables. Strong positive and negative correlations between features are observed. For example, the number of non-H atoms and number of non-C/H atoms are strongly correlated positively in both the training set and the test set. For another example, the molecular weight is strongly correlated with Gaussian1, Gaussian2, total surface area, molecular complexity, and non-H atoms.

**Fig. 8.**
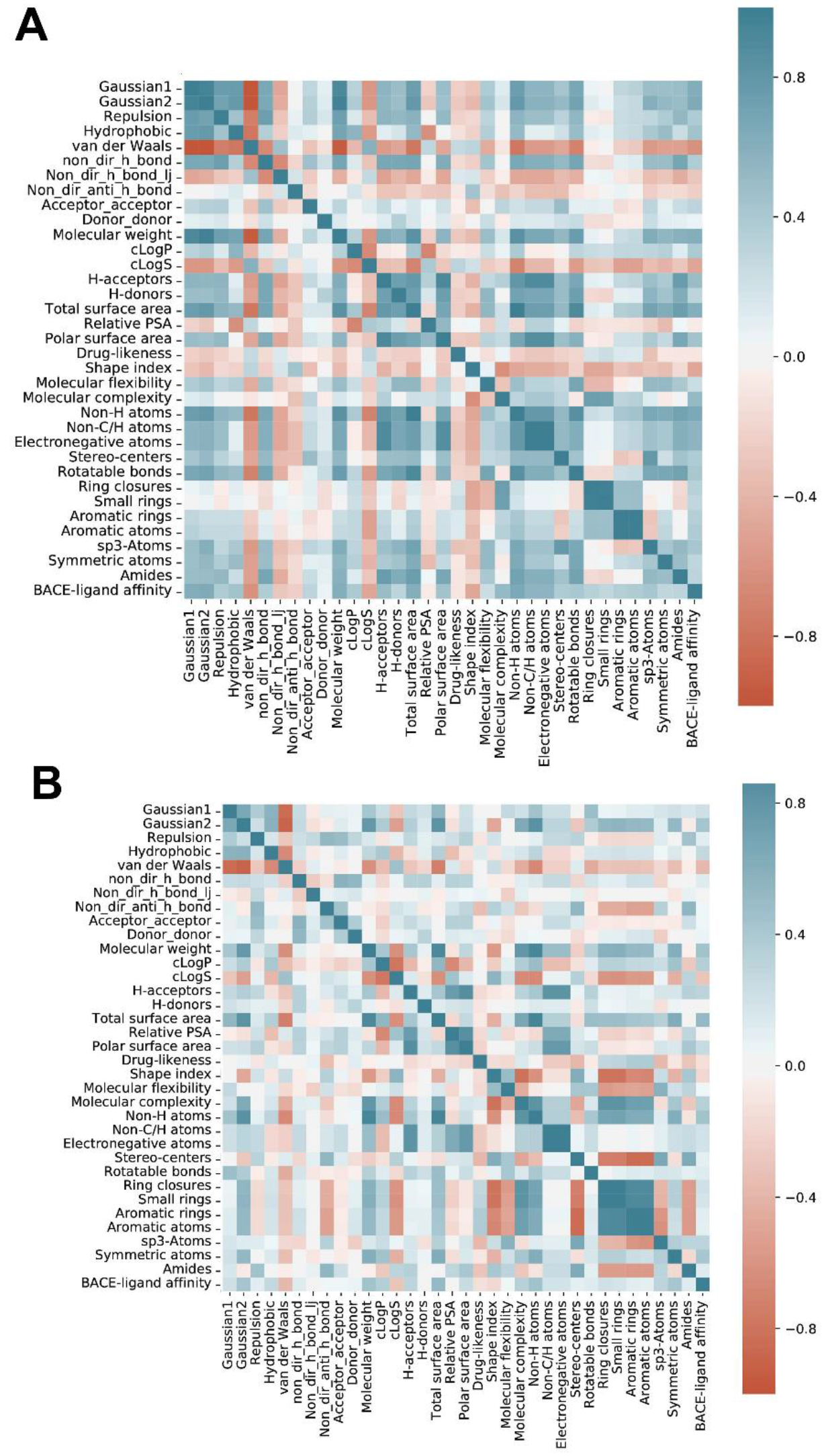
Normalized covariance matrix of features in the training set and D3R GC4 BACE test set. A, covariance matrix of the 34 features against the BACE training set (sample size: 222). B, covariance matrix of the 34 features against the D3R GC4 BACE test set (sample size: 154). The variance of each feature is scaled to unity using the corrcoef function in Numpy. Matplotlib was also used to generate this figure.

To better understand our MLP(10,8,8) model using structure and ligand-based features, a feature importance analysis was performed on the BACE training set. To estimate the feature importance of each feature, one feature at a time was set to the mean of this feature before the 10-fold crossvalidation. The means of R^2^ in the 10-fold validations and standard deviations for each feature were recorded, are compiled in **Table 9** and shown in **Fig. 9A**. The difference of model performance (using R^2^) before and after setting to mean of each feature is considered the feature importance. Interestingly, the top three important features evaluated using all 34 features are Gaussian1, clogP, and number of rotatable bonds, which are not the strongest affinity correlated features in the training set, namely: Gaussian2, van der Waals, molecular weight, total surface area, and number of non-H. This discrepancy is probably because the features with high importance already contain the information from the top affinity correlated features. For example, Gaussian1 correlates well with Gaussian2 and molecular weight. Thus, even though Gaussian2 and molecular weight show less importance in model performance, Gaussian1 already conveys the ranking power originated from Gaussian2 and molecular weight.

**Fig 9.**
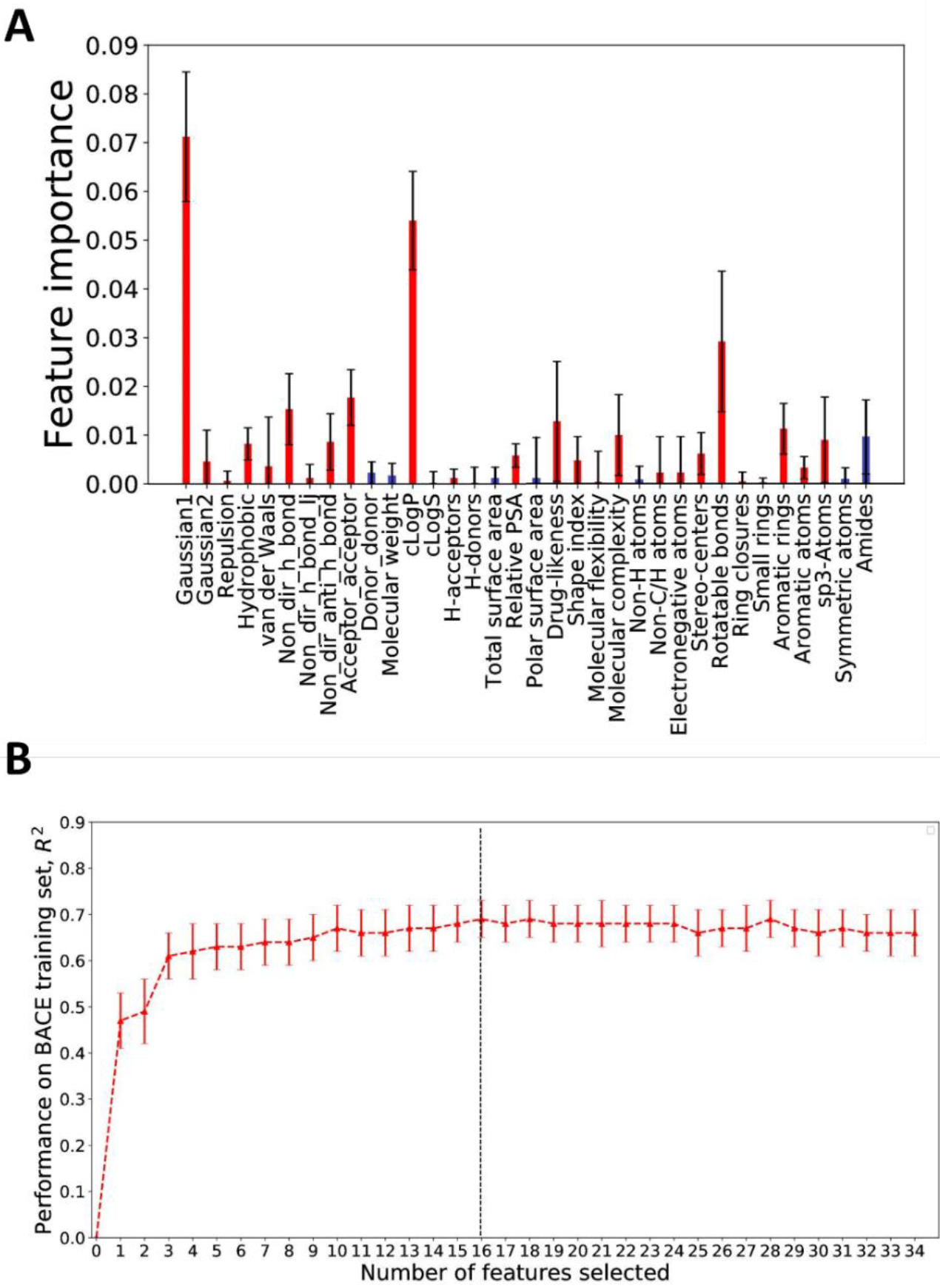
Feature importance and recursive feature elimination analysis of the D3R submitted MLP(10,8,8) model using 34 structure- and ligand-based features against the BACE training set. A, feature importance analysis of the MLP model, reflected by the performance difference in R^2^. The differences in performance before and after setting each feature to its mean was evaluated in 10-fold cross-validation using the BACE training set (sample size = 154). Red rectangles indicate that muting that feature lowers the model performance (indicating a positive contribution of this feature). Blue rectangles indicate that muting that feature enhances the model performance (indicating the feature only contributes noise). The error bar is the standard error in the 10-fold cross-validation. B, recursive feature elimination analysis of the MLP model. The error bar is the standard error in the 10-fold validation. The dashed line at 16 features indicate the optimal performance using this methodology.

**Table 9.** Feature importance, importance rank and sequence in recursive feature elimination(RFE) in the D3R submitted MLP(10,8,8) model against the BACE training set (sample size = 222). Lower RFE sequence numbers indicates weaker feature contributions towards the overall model performance.

|  | Features | Feature importance | Feature importance rank | Sequence in RFE |
| --- | --- | --- | --- | --- |
| 1 | Gaussian1 | 0.071 | 1 | 34 |
| 2 | Gaussian2 | 0.005 | 15 | 11 |
| 3 | Repulsion | 0.001 | 22 | 21 |
| 4 | Hydrophobic | 0.008 | 11 | 33 |
| 5 | van der Waals | 0.004 | 16 | 29 |
| 6 | Non_dir_h_bond | 0.015 | 5 | 7 |
| 7 | Non_dir_h_bond_lj | 0.001 | 21 | 3 |
| 8 | Non_dir_anti_h_bond | 0.009 | 10 | 2 |
| 9 | Acceptor_acceptor | 0.018 | 4 | 13 |
| 10 | Donor_donor | -0.002 | 33 | 14 |
| 11 | Molecular weight | -0.002 | 32 | 10 |
| 12 | cLogP | 0.054 | 2 | 31 |
| 13 | cLogS | 0.000 | 26 | 28 |
| 14 | H-acceptors | 0.001 | 20 | 15 |
| 15 | H-donors | 0.000 | 27 | 23 |
| 16 | Total surface area | -0.001 | 30 | 20 |
| 17 | Relative PSA | 0.006 | 13 | 16 |
| 18 | Polar surface area | -0.001 | 31 | 17 |
| 19 | Drug-likeness | 0.013 | 6 | 26 |
| 20 | Shape index | 0.005 | 14 | 32 |
| 21 | Molecular flexibility | 0.000 | 24 | 30 |
| 22 | Molecular complexity | 0.010 | 8 | 8 |
| 23 | Non-H atoms | -0.001 | 28 | 25 |
| 24 | Non-C/H atoms | 0.002 | 19 | 5 |
| 25 | Electronegative atoms | 0.002 | 18 | 19 |
| 26 | Stereo-centers | 0.006 | 12 | 6 |
| 27 | Rotatable bonds | 0.029 | 3 | 9 |
| 28 | Ring closures | 0.000 | 23 | 18 |
| 29 | Small rings | 0.000 | 25 | 4 |
| 30 | Aromatic rings | 0.011 | 7 | 27 |
| 31 | Aromatic atoms | 0.003 | 17 | 24 |
| 32 | sp3-Atoms | 0.009 | 9 | 12 |
| 33 | Symmetric atoms | -0.001 | 29 | 22 |
| 34 | Amide | -0.010 | 34 | 1 |

\*R<sup>2</sup> reduction after the feature is set to the mean in the 10-fold cross-validated BACE training set.

Recursive feature elimination (RFE), where the weakest feature is removed in each iteration until a satisfactory performance is achieved, was conducted. The result is shown in **Fig. 9B**. The sequence of feature removal is shown in **Table 9**. In general, features with less importance tend to be eliminated first. For example, the weakest feature, the number of amide groups, is removed first. However, apparent variations on the feature removal sequence against the ranking of feature importance are observed. Note the last three features left in RFE are: Gaussian1, hydrophobic term, and shape index, compared to Gaussian1, clogP, and number of rotatable bonds in feature importance analysis. This shows that the features remaining in RFE are not necessarily those with the highest feature importance, primarily due to the correlation between features.

Due to time constraints, we did not analyze the features before our submission of the MLP model. Here, we compare the performance with/without the RFE. According to this RFE analysis, when the top 16 features (after 18 features were removed) are used, the MLP(10,8,8) model (λ = 10) has a slightly improved prediction ability, with a R^2^ of 0.69(12) and 0.755(9) to the training and test sets in 10-fold cross-validations in X_training_. Using the selected 16 features and same architecture of the MLP(10,8,8) model, we adjusted the regularization parameter (λ). When λ is in range of 5-10, the MLP model has a R^2^ of 0.69(14) against the 10-fold cross-validation training set. However, the model gained no improvement toward the D3R test set, as the Spearman’s p is 0.41, 0.42, 0.43 for λ at 5, 7, 10.

In summary, the feature importance analysis and recursive feature elimination, provided a slight improvement for the MLP model against the BACE training set but did generate an improved model against the D3R blinded test set.

## Conclusions

Through participating in the D3R GC4 BACE Subchallenge, we investigated optimizing the AutoDock Vina docking scoring function and designed and compared machine learning models using a combination of structure-based and ligand-based terms. To generate docked poses for BACE macrocyclic ligands, a 3D similarity pose automated selection script was shown to be effective in generating accurate docked poses. Five different machine learning models were explored ranging in complexity from linear regression to a deep neural network. From tuning different models, we found that hyperparameter tuning greatly affects the accuracy of protein-ligand affinity prediction. Feature importance and recursive feature elimination analysis have also been shown to be useful in improving model performance against the compiled training set. However, incorporating the feature analysis retrospectively does not enhance the model performance on the D3R blind set.

This work shows that non-linear machine learning models are highly effective for protein-ligand affinity prediction if high-quality training datasets are available for the target protein. We found that the Vinardo scoring function, developed from a broad set of ligands, performed best for docking macrocyclic ligands to BACE. We expect docking performance can be further improved by careful selection and optimization of receptor structures as well as extensive ligand conformer generation. In contrast, a deep neural network trained specifically on BACE ligands performed best for affinity prediction. However, due to the similarity of the macrocyclic ligands in the D3R GC4 contest, the feature extraction methods we utilized need more consideration in order to improve the affinity prediction power. Affinity prediction can probably be improved by training on larger datasets, training on ligand/target-specific datasets, using deeper neural networks and adopting advanced neural networks such as convolutional neural networks, automated tuning of hyperparameters, including chemical topological features, and carefully selecting a larger set of informative input features.

## Acknowledgments

This work was funded by NSF CAREER MCB 1833181.

