## Supporting Tables 1-2 for "Deep neural network affinity model for BACE inhibitors in D3R Grand Challenge 4"

Bo Wang · Ho-Leung Ng

**Table S1** Pairing of test ligands with released ligands……………………………………………….….. *p.* 2

**Table S2** BACE affinity dataset.………………………………………..……………………….. *pp.* 3-4

**Table S1.** Pairing of test ligands with released ligands. The FragFp method was used in Data Warrior.

| Test ligands | Released ligands | Test ligands | Released ligands | Test ligands | Released ligands | Test ligands | Released ligands |
|---|---|---|---|---|---|---|---|
| BACE_1 | BACE_1 | BACE_135 | BACE_16 | BACE_32 | BACE_3 | BACE_69 | BACE_9 |
| BACE_10 | BACE_10 | BACE_136 | BACE_15 | BACE_33 | BACE_3 | BACE_7 | BACE_7 |
| BACE_100 | BACE_15 | BACE_137 | BACE_12 | BACE_34 | BACE_4 | BACE_70 | BACE_8 |
| BACE_101 | BACE_15 | BACE_138 | BACE_14 | BACE_35 | BACE_4 | BACE_71 | BACE_9 |
| BACE_102 | BACE_14 | BACE_139 | BACE_15 | BACE_36 | BACE_2 | BACE_72 | BACE_10 |
| BACE_103 | BACE_12 | BACE_14 | BACE_14 | BACE_37 | BACE_3 | BACE_73 | BACE_10 |
| BACE_104 | BACE_17 | BACE_140 | BACE_14 | BACE_38 | BACE_19 | BACE_74 | BACE_8 |
| BACE_105 | BACE_15 | BACE_141 | BACE_15 | BACE_39 | BACE_5 | BACE_75 | BACE_10 |
| BACE_106 | BACE_14 | BACE_142 | BACE_10 | BACE_4 | BACE_4 | BACE_76 | BACE_10 |
| BACE_107 | BACE_14 | BACE_143 | BACE_3 | BACE_40 | BACE_7 | BACE_77 | BACE_14 |
| BACE_108 | BACE_14 | BACE_144 | BACE_14 | BACE_41 | BACE_8 | BACE_78 | BACE_18 |
| BACE_109 | BACE_14 | BACE_145 | BACE_14 | BACE_42 | BACE_8 | BACE_79 | BACE_18 |
| BACE_11 | BACE_11 | BACE_146 | BACE_15 | BACE_43 | BACE_8 | BACE_8 | BACE_8 |
| BACE_110 | BACE_14 | BACE_147 | BACE_19 | BACE_44 | BACE_8 | BACE_80 | BACE_9 |
| BACE_111 | BACE_14 | BACE_148 | BACE_3 | BACE_45 | BACE_12 | BACE_81 | BACE_14 |
| BACE_112 | BACE_14 | BACE_149 | BACE_3 | BACE_46 | BACE_8 | BACE_82 | BACE_9 |
| BACE_113 | BACE_12 | BACE_15 | BACE_15 | BACE_47 | BACE_8 | BACE_83 | BACE_18 |
| BACE_114 | BACE_14 | BACE_150 | BACE_19 | BACE_48 | BACE_8 | BACE_84 | BACE_8 |
| BACE_115 | BACE_14 | BACE_151 | BACE_19 | BACE_49 | BACE_12 | BACE_85 | BACE_3 |
| BACE_116 | BACE_14 | BACE_152 | BACE_20 | BACE_5 | BACE_5 | BACE_86 | BACE_3 |
| BACE_117 | BACE_14 | BACE_153 | BACE_20 | BACE_50 | BACE_6 | BACE_87 | BACE_11 |
| BACE_118 | BACE_12 | BACE_154 | BACE_20 | BACE_51 | BACE_8 | BACE_88 | BACE_11 |
| BACE_119 | BACE_14 | BACE_155 | BACE_20 | BACE_52 | BACE_8 | BACE_89 | BACE_11 |
| BACE_12 | BACE_12 | BACE_156 | BACE_19 | BACE_53 | BACE_9 | BACE_9 | BACE_9 |
| BACE_120 | BACE_14 | BACE_157 | BACE_20 | BACE_54 | BACE_8 | BACE_90 | BACE_11 |
| BACE_121 | BACE_14 | BACE_158 | BACE_20 | BACE_55 | BACE_9 | BACE_91 | BACE_2 |
| BACE_122 | BACE_14 | BACE_16 | BACE_16 | BACE_56 | BACE_10 | BACE_92 | BACE_11 |
| BACE_123 | BACE_14 | BACE_19 | BACE_19 | BACE_57 | BACE_10 | BACE_93 | BACE_11 |
| BACE_124 | BACE_14 | BACE_20 | BACE_20 | BACE_58 | BACE_9 | BACE_94 | BACE_11 |
| BACE_125 | BACE_14 | BACE_21 | BACE_3 | BACE_59 | BACE_9 | BACE_95 | BACE_14 |
| BACE_126 | BACE_17 | BACE_22 | BACE_3 | BACE_6 | BACE_6 | BACE_96 | BACE_14 |
| BACE_127 | BACE_15 | BACE_23 | BACE_4 | BACE_60 | BACE_10 | BACE_97 | BACE_12 |
| BACE_128 | BACE_14 | BACE_24 | BACE_2 | BACE_61 | BACE_9 | BACE_98 | BACE_14 |
| BACE_129 | BACE_15 | BACE_25 | BACE_3 | BACE_62 | BACE_9 | BACE_99 | BACE_12 |
| BACE_13 | BACE_13 | BACE_26 | BACE_3 | BACE_63 | BACE_2 | | |
| BACE_130 | BACE_15 | BACE_27 | BACE_5 | BACE_64 | BACE_9 | | |
| BACE_131 | BACE_14 | BACE_28 | BACE_3 | BACE_65 | BACE_9 | | |
| BACE_132 | BACE_15 | BACE_29 | BACE_5 | BACE_66 | BACE_9 | | |
| BACE_133 | BACE_14 | BACE_30 | BACE_5 | BACE_67 | BACE_10 | | |
| BACE_134 | BACE_15 | BACE_31 | BACE_5 | BACE_68 | BACE_9 | | |

**Table S2**. BACE affinity training dataset. It contains 222 BACE-ligand affinities extracted from PDBbind set 2017. The affinity is estimated by -log($K_i$, /$K_d$, or /$IC_{50}$), where $K_i$, $K_d$, and $IC_{50}$ is in mol/L unit.

| PDB | Ligand | Affinity | PDB | Ligand | Affinity | PDB | Ligand | Affinity |
|---|---|---|---|---|---|---|---|---|
| 1W51 | L01 | 6.30 | 3NSH | 957 | 6.33 | 4HA5 | 13W | 7.24 |
| 1XS7 | MMI | 7.60 | 3OOZ | ZOO | 7.85 | 4HZT | 0ZA | 6.12 |
| 2F3E | AXQ | 6.81 | 3PI5 | 3P5 | 6.02 | 4I0D | 1B7 | 6.21 |
| 2F3F | AXF | 6.72 | 3QBH | QBH | 6.82 | 4I0F | 1BF | 6.35 |
| 2FDP | FRP | 7.59 | 3R2F | PB0 | 8.85 | 4I0Z | 1BB | 6.33 |
| 2G94 | ZPQ | 9.52 | 3RSV | 3RS | 9.15 | 4I10 | 1BS | 6.77 |
| 2HIZ | LIJ | 6.95 | 3RSX | RSV | 4.41 | 4I11 | 1CH | 4.57 |
| 2HM1 | LIQ | 8.70 | 3RTH | RTH | 5.21 | 4I12 | 1BC | 6.39 |
| 2IQG | F2I | 8.30 | 3RTM | RTM | 4.42 | 4I1C | 1BE | 8.10 |
| 2OHK | 1SQ | 2.70 | 3RTN | RTN | 7.13 | 4JOO | 1M4 | 4.43 |
| 2OHL | 2AQ | 3.05 | 3RU1 | 3RU | 4.85 | 4JP9 | 1M5 | 7.62 |
| 2OHM | 8AP | 3.51 | 3RVI | RVI | 7.96 | 4JPC | 1M6 | 7.03 |
| 2OHP | 6IP | 4.03 | 3S2O | EV6 | 5.05 | 4JPE | 1M7 | 7.32 |
| 2OHQ | 7IP | 4.60 | 3S7L | 591 | 7.10 | 4K8S | 1QT | 7.44 |
| 2OHR | 8IP | 4.00 | 3S7M | 532 | 8.00 | 4K9H | 1QU | 7.12 |
| 2OHS | 9IP | 4.40 | 3SKF | PB7 | 8.30 | 4KE0 | 1R8 | 7.77 |
| 2OHT | IP6 | 5.04 | 3SKG | PB8 | 8.10 | 4KE1 | 1R6 | 8.60 |
| 2OHU | IP7 | 5.38 | 3TPR | 5HA | 7.70 | 4L7G | 1W0 | 4.00 |
| 2P4J | 23I | 8.96 | 3U6A | 18P | 6.49 | 4L7H | 1W1 | 3.70 |
| 2P83 | MR0 | 7.96 | 3UDH | 91 | 2.85 | 4L7J | 1W2 | 3.70 |
| 2QK5 | CS5 | 8.10 | 3UDJ | 92 | 3.62 | 4LC7 | 1WP | 4.93 |
| 2QMD | CS7 | 8.30 | 3UDK | 95 | 4.49 | 4LXA | 1YS | 8.70 |
| 2QMF | CS9 | 8.52 | 3UDM | 09A | 4.48 | 4LXK | 1YT | 8.40 |
| 2QMG | SC6 | 9.15 | 3UDN | 09B | 5.15 | 4LXM | 1YU | 7.06 |
| 2QP8 | SC7 | 8.10 | 3UDP | 09D | 5.22 | 4N00 | 2EX | 8.02 |
| 2QU2 | 251 | 5.43 | 3UDQ | 09E | 6.00 | 4PZW | 2X4 | 6.96 |
| 2QU3 | 462 | 6.23 | 3UDR | 09F | 4.96 | 4PZX | 2X5 | 6.00 |
| 2VKM | BSD | 8.74 | 3UDY | 09G | 5.70 | 4R5N | 3J9 | 7.12 |
| 2ZDZ | 310 | 6.15 | 3UFL | 508 | 6.82 | 4R8Y | 3KO | 6.19 |
| 2ZE1 | 411 | 6.22 | 3VEU | 0GO | 7.26 | 4R91 | 3KT | 6.37 |
| 3CIB | 314 | 7.85 | 3VF3 | 0GS | 5.86 | 4R92 | 3KU | 7.41 |
| 3CIC | 316 | 8.52 | 3VG1 | 0GT | 7.26 | 4R93 | 779 | 7.80 |
| 3CID | 318 | 8.30 | 3VV6 | B00 | 3.80 | 4R95 | 3KW | 7.85 |
| 3CKP | 12 | 6.35 | 3WB4 | 0B3 | 4.44 | 4RCD | 3LL | 9.15 |
| 3CKR | 9 | 5.30 | 3WB5 | 0B4 | 4.57 | 4RCE | 3LN | 8.70 |
| 3DM6 | 757 | 7.43 | 4ACU | QN7 | 7.39 | 4RCF | 3LO | 8.40 |
| 3DUY | AFJ | 5.85 | 4ACX | S8Z | 7.11 | 4RRN | 3UW | 7.32 |

**Formatted:** Subscript
**Formatted:** Subscript
**Formatted:** Subscript
**Formatted:** Subscript
**Formatted:** Subscript
**Formatted:** Subscript

| | | | | | | | | |
|---|---|---|---|---|---|---|---|---|
| 3DV1 | AR9 | 6.23 | 4B1C | 1B1 | 7.10 | 4RRO | 3UX | 7.59 |
| 3DV5 | BAV | 7.66 | 4B1D | 6TG | 7.49 | 4RRS | 3UY | 6.68 |
| 3H0B | B35 | 5.49 | 4B70 | WM9 | 5.20 | 4WTU | 3UT | 9.52 |
| 3HVG | EV0 | 2.70 | 4B72 | 2FB | 6.20 | 4WY1 | 3VO | 5.29 |
| 3HW1 | EV2 | 3.06 | 4B77 | 54M | 5.10 | 4WY6 | 3VP | 5.97 |
| 3I25 | MV7 | 8.51 | 4B78 | KGG | 5.00 | 4X2L | 3WP | 4.44 |
| 3IGB | 454 | 4.42 | 4D83 | 0GT | 7.26 | 4X7I | 3YS | 7.69 |
| 3IN3 | 472 | 7.22 | 4D85 | 0GU | 6.85 | 4XKX | 43K | 9.05 |
| 3IN4 | BX2 | 7.52 | 4D88 | BXQ | 7.31 | 4XXS | SI5 | 7.28 |
| 3IND | 593 | 5.82 | 4D89 | BXD | 8.70 | 4YBI | 4B2 | 6.60 |
| 3INE | X17 | 6.77 | 4D8C | BXD | 8.70 | 4ZSM | 4RW | 2.40 |
| 3INF | X45 | 7.40 | 4DH6 | 0KN | 7.77 | 4ZSP | 4RZ | 5.74 |
| 3INH | 569 | 7.70 | 4DI2 | 0K9 | 8.26 | 4ZSQ | 4RX | 5.65 |
| 3IVH | 1LI | 7.33 | 4DJU | 0KK | 5.44 | 4ZSR | 4RY | 4.44 |
| 3IVI | 2LI | 7.92 | 4DJV | 0KM | 6.72 | 5CLM | 52K | 7.36 |
| 3IXJ | 586 | 9.49 | 4DJW | 0KP | 6.28 | 5DQC | 5E6 | 7.34 |
| 3IXK | 929 | 8.18 | 4DJX | 0KQ | 7.23 | 5ENK | 5QV | 8.10 |
| 3K5C | 0BI | 7.77 | 4DJY | 0KR | 8.27 | 5ENM | 5QU | 7.15 |
| 3K5D | XLI | 7.28 | 4DPF | 0LG | 6.40 | 5HD0 | 60Y | 8.40 |
| 3K5F | AYH | 5.43 | 4DPI | 0N1 | 7.11 | 5HDU | 60W | 9.00 |
| 3K5G | BJC | 8.60 | 4DUS | 0MP | 8.30 | 5HDV | 60V | 7.82 |
| 3KYR | 38 | 6.72 | 4EWO | 996 | 7.36 | 5HDX | 60U | 7.32 |
| 3L38 | 879 | 7.00 | 4EXG | 916 | 8.07 | 5HDZ | 954 | 7.35 |
| 3L3A | 625 | 6.38 | 4FM7 | 0UP | 7.24 | 5HE4 | 60T | 7.70 |
| 3L58 | CS5 | 8.10 | 4FM8 | 0UQ | 5.97 | 5HE5 | 60S | 6.47 |
| 3L59 | BDJ | 3.70 | 4FRI | DWA | 5.55 | 5HE7 | 60X | 8.30 |
| 3L5B | BDO | 3.91 | 4FRJ | DWB | 6.55 | 5HTZ | 66J | 8.62 |
| 3L5C | BDQ | 5.15 | 4FRK | DWD | 8.10 | 5HU0 | 66H | 6.23 |
| 3L5D | BDV | 4.11 | 4FRS | 0V6 | 8.77 | 5HU1 | 66F | 8.66 |
| 3L5E | BDW | 7.57 | 4FSE | 0VA | 7.40 | 5I3V | 68M | 7.80 |
| 3L5F | BDX | 6.22 | 4FSL | 0VB | 7.70 | 5I3W | 68L | 9.22 |
| 3LHG | Z81 | 7.70 | 4GID | 0GH | 10.77 | 5I3X | 68J | 8.10 |
| 3LNK | 74A | 5.94 | 4H1E | 10J | 8.52 | 5I3Y | 68K | 9.40 |
| 3MSJ | EV3 | 3.11 | 4H3F | 10O | 9.00 | 5IE1 | 6BS | 6.85 |
| 3MSK | EV4 | 4.59 | 4H3G | 10Q | 8.22 | 5KQF | 6WD | 6.51 |
| 3MSL | EV5 | 5.15 | 4H3I | 10V | 8.52 | 5KR8 | 6WE | 6.82 |
| 3N4L | 842 | 7.23 | 4H3J | 10W | 7.05 | 5TOL | 7H3 | 7.48 |
